# 3D chromatin maps of a brown alga reveal U/V sex chromosome spatial organization

**DOI:** 10.1101/2024.05.11.593484

**Authors:** Pengfei Liu, Jeromine Vigneau, Rory Craig, Josue Barrera-Redondo, Elena Avdievich, Claudia Martinho, Michael Borg, Fabian B. Haas, Chang Liu, Susana M Coelho

## Abstract

Sex chromosomes are unique genomic regions displaying structural and evolutionary features that distinguish them markedly from autosomes. Although nuclear three dimensional (3D) folding of chromatin structure is important for gene expression regulation and correct developmental programs, very little is known about the 3D architecture of sex chromosomes within the nucleus, and how that impacts their function in sex determination. Here, we determine the sex-specific 3D organization of the model brown alga *Ectocarpus* chromosomes at 2 kb resolution, by comprehensively mapping long-range chromosomal interactions using Hi-C coupled with Oxford Nanopore long reads. We report that *Ectocarpus* interphase chromatin exhibits a non-Rabl conformation, with strong contacts among telomeres and among centromeres, which feature centromere-specific LTR retrotransposons. The *Ectocarpus* chromosomes do not contain large local interactive domains that resemble TADs described in animals, but their 3D genome organization is largely shaped by post-translational modifications of histone proteins that regulate chromatin compaction and mediate transcriptional regulation. We describe the spatial conformation and sub-nuclear positioning of the sex determining region (SDR) within the U and V chromosomes and show that these regions are very insulated and span the centromeres. Moreover, we link sex-specific chromatin dynamics and gene expression levels to the 3D chromatin structure of U and V chromosomes. Finally, we uncover the unique conformation of a large genomic region on chromosome 6 harboring an endogenous viral element (EVE), providing insights regarding the functional significance of the chromatin organisation of latent giant dsDNA virus.

## Introduction

Sex chromosomes are unique genomic regions that evolved independently many times in different groups of eukaryotes. Three types of sex chromosome system exist in nature, the well-described XX/XY and ZW/ZZ systems, and the still elusive UV systems, in organisms that express sex in the haploid stage of the life cycle ^1–6^. Heteromorphic sex chromosomes (Y, W, U, V) have evolved repeatedly in diverse eukaryotic species. Suppression of recombination between X and Y (or Z and W, or U and V) chromosomes usually leads to a range of genomic modifications in this regions, including degeneration of the non-recombining chromosome, accumulation of repeats, and gene decay via accumulation of deleterious mutations ^7,8^. Repeats pose the largest challenge for reference genome assembly, and centromeres, subtelomeres and the repeat-rich sex chromosome are typically ignored from se-quencing projects. Consequently, complete sequence assemblies of heteromorphic Y, W, U and V sex chromosomes have only been generated across a handful of taxa ^9–16^, and most of the information is fragmentary even at the linear sequence level. Moreover, despite the key role that the 3D structure of chromatin plays in gene regulation ^17,18^, we lack critical information regarding chromatin landscapes and nuclear 3D organization of sex chromosomes within the nuclear space, and how chromatin folding is associated with the sex-specific gene expression underlying sexual differentiation.

Genome folding generally involves hierarchical structures ranging from chromatin loops to chromosome territories ^19^. The best-known 3D chromatin organization units are topologically associating domains (TADs), which show a self-interacting pattern with strongly interacting boundaries in Hi-C contact maps of animal genomes ^20^. Genome architectural proteins, such as CTCF (CCCTC-binding factor) and cohesin, bind strongly to DNA anchor sites and mediate the formation of chromatin contact domains through loop extrusion ^21^. In addition to TADs, structural units called compartmental domains have been demonstrated in animals (e.g. ^22^). Compartmental domains are closely associated with local chromatin states and preferentially interact with other compartmental domains of similar chromatin states, contributing to the establishment of the 3D architecture for a given genome ^19,22^. Plant genomes also frequently exhibit a higher-order 3D chromatin organization. TADs or TAD-like structures have been described in several plant species ^23^, although their genomes do not encode CTCF homologs ^24^. In contrast, *Arabidopsis* (*Arabidopsis thaliana*) lacks plant-type TADs ^25^. The absence of TADs in the *Arabidopsis* genome is likely related to its small size, high gene density and short intergenic regions. However chromatin loops and compartments are present in *A. thaliana* (e.g. ^26^) and small structural units within 3D chromatin architecture have been recently described ^27^.

Here, we generated 2 kb-resolution maps of the male and female haploid genomes of the brown algal model *Ectocarpus*, and examined the 3D chromatin structure of autosomes compared to U and V sex chromosomes. *Ectocarpus* life cycle involves an alternation between diploid and haploid generations, with sex being determined in the haploid stage of the life cycle by U (female) and V (male) sex chromosomes ^5^. Therefore, this model organism provides the opportunity to investigate the sub-nuclear organization of the chromatin structure of U/V sex chromosomes and compare it to autosomes. Our near complete assembly of the *Ectocarpus* genome (*Ectocarpus* V5) offers an improved reference genome and allowed us to define and characterise the centromeric and sub-telomeric sequences in this organism. We found that interphase chromatin is organized in a non-Rabl configuration, with telomeres and centromeres of all 27 *Ectocarpus* chromosomes clustering together in the 3D nuclear space. We reveal that the 3D structure of *Ectocarpus* chromatin is highly streamlined, not organized into TADs, and A and B chromatin compartments are mainly defined by H3K79me2 deposition and depletion of activation marks. We then focus on the 3D structure of the *Ectocarpus* U and V sex chromosomes to show that in both sex chromosomes the SDR spans the centromere, and is highly insulated from the rest of the chromosome. We found no overall differences in the 3D chromatin organization between male (V) and female (U) chromosomes but both have different 3D organization compared with autosomes. Finally, we uncover the distinctive conformation of a genomic region on chromosome 6 harboring a giant endogenous viral element (EVE), giving insights into the interplay of dsDNA viruses with the chromatin environment in the host.

## Results

### A near complete assembly of the male and female haploid genome of *Ectocarpus*

A complete assembly of the *Ectocarpus* genome has been challenging mainly due to the presence of highly repetitive regions, which short-read Illumina sequencing, low coverage Hi-C and Sanger sequencing could not hitherto successfully resolve. The published version of the *Ectocarpus* sp.7 reference genome (V2) contains 28 pseudo-chromosomes spanning 176.99 Mb, with 17.97 Mb of unplaced contigs, a contig-level N50 of 33 kb and a total of 11,588 gaps ^28^. Here, we combined Oxford Nanopore Technologies (ONT) long reads and Hi*-*C sequencing techniques to achieve near complete assemblies of both the haploid male and female genome of *Ectocarpus* sp.7 (Figure 1A, Table S1-S2).

**Figure 1.**
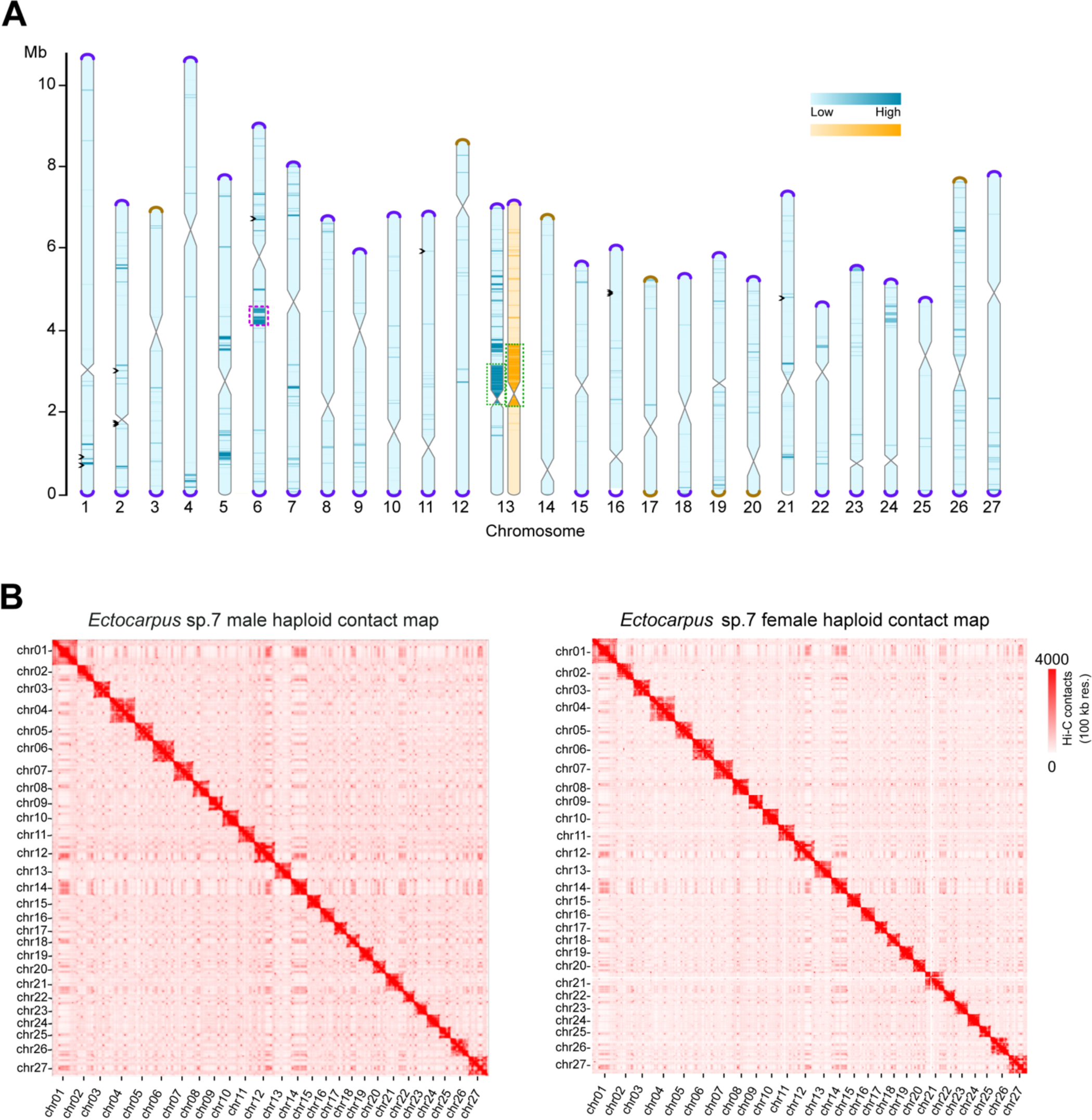
Ectocarpus sp7 whole-genome assembly. (A) Schematic representation of the near telomere-to-telomere assembly of the 27 Ectocarpus sp.7 chromosomes, in haploid male (blue) and female (orange). Telomeres are represented as violet caps, sub-telomeres in brown. Centromeric regions are represented by the constrictions in the center of the chromosomes. The chromosomes are filled by variant density between the male and female haploid genomes used for the assembly (darker color means more differences). Violet dotted boxes represent the genomic region where a dsDNA virus is inserted and green dotted boxes represent the SDRs. Black arrowheads depict gaps. See methods for details. (B) Genome-wide Hi-C contact map showing frequencies of pairwise 3D genome contacts at a 100kb resolution in the male and female haploid genomes.

The ONT long reads were obtained separately from male and female siblings (Ec32m, Ec25f, Figure S1, S2), totaling 11 Gb and 20 Gb of data, respectively. ONT long-reads were complemented with Hi-C data, encompassing 822 million pairs of sequences (∼135 Gb) at a sequencing depth of 635x for the male, and ∼444 million pairs (∼73 Gb) and 338x coverage for the female (see methods for details) (Figure S1, Table S3). The *Ectocarpus* V5 male genome assembly has an N50 of 7.0 Mb and a total size of 186.6 Mb. The V2 chromosome 28 from V2 is now part of chromosome 4, bringing the total number of chromosomes to 27, with sizes ranging from 4.52 Mb to 10.73 Mb. Similarly to what was done for the *Ectocarpus* V2 ^28^, we added the female SDR (size 1.55 Mb) to this male genome in order to obtain a final *Ectocarpus* V5 reference genome (Figure S1).

The genome is highly contiguous: out of the 27 assembled chromosome models, the majority contain zero gaps; and six chromosomes have only 10 gaps in total (Figure 1A, Table S4). The accuracy of Hi-C based chromosome construction was evaluated manually by inspecting the chromatin contact matrix at 100 kb resolution, which exhibited a well-organized interaction contact pattern along the diagonals within each pseudo-chromosome (Figure 1B).

Telomeric regions were almost entirely absent from earlier genome assemblies, although a putative telomere bearing the repeat (TTAGGG)n was observed ^29^. In our *Ectocarpus* V5 assembly, 43 of the 54 telomeric regions are fully resolved, and twelve of the 27 pseudo-chromosomes correspond to a telomere-to-telomere assembly (Figure 1A, Figure S3). On all but three of the resolved telomeric regions, we observed a specific satellite repeat adjacent to the telomeric repeats. The satellite features a repeated monomer of ∼98 bp and is almost exclusively found at the sub-telomeres, where it forms arrays that range from only a few to more than 100 copies. Notably, the telomeric motif TTAGGG is present in three independent locations with each satellite monomer (Figure S4). Similar subtelomeric organizations have been observed in several species including the green alga *Chlamydomonas reinhardtii*, where the telomere-like motifs present within the sub-telomeric satellites are hypothesised to serve as seed sequences that facilitate telomere healing following DNA damage ^30^. Eight of the V5 chromosome arms terminated in the subtelomeric repeat, leaving only four chromosome extremities for which the assembly failed to reach either the subtelomere or telomere (Figure 1A).

Ribosomal DNA (rDNA) arrays were also poorly resolved in previous assemblies. The V5 assembly revealed a single major rDNA array located within an internal region of chromosome 4, which features six rDNA repeats before resulting in a contig break. The 5S rDNA gene is linked to the main rDNA unit (18S-5.8S-26S), as previously reported in many brown algae and Stramenopiles^31^. Considering ONT read coverage of 7,000x, we estimate that the rDNA array may consist of >100 copies, spanning ∼800 kb.

The total repeat content of the assembled chromosomes is estimated to be 29.8% (Table S4). 13.75 Mb of addi-tional sequence could not be assembled to chromosomes and remains unplaced in the V5 assembly. These se-quences are extremely repetitive (74.3% repeats) and presumably include heterochromatic regions that corre-spond to some of the assembly gaps or incomplete chromosome ends. Longer reads or alternative technologies will be required to achieve complete assembly of these complex regions.

Since the V2 genome had a high-quality gene annotation, we performed a lift over of the V2 gene models to the *Ectocarpus* V5 genome. Out of the 18,412 V2 gene models, 18,278 could be lifted, and the remaining were mostly located on an unassigned scaffold in the V2 assembly. Genome completeness was quantified by BUSCO ^32^. Two database sets were used, eukaryota (255 core genes) and stramenopiles (100 core genes). Of the 255 core eukar-yotic BUSCO genes, the V5 reference assembly contains 226 (88.7%) complete BUSCO genes. This represents a gain of 8 genes (+3.2%) compared to the V2 genome. The stramenopiles result increased by 1 % (Table S5).

### The *Ectocarpus* 3D chromatin architecture

To explore the 3D chromatin architecture of *Ectocarpus*, we mapped male and female Hi-C reads back to the V5 assembly (Table S3, Figure S1). Biological replicates were highly correlated (Pearson r= 0.96 and r=0.94 for male and female samples, respectively, Figure S5), therefore, replicates were combined for downstream analysis to produce sex-specific high-resolution maps. We obtained 188.8 and 134.8 million interaction read pairs for male and female, respectively, reaching a 2 kb resolution for each of the sexes.

In animals and plants, chromosomes are hierarchically packed in the nuclear space, and each occupies discrete regions referred to as a chromosome territory (CT) ^23,33^. Chromosomal territories were detected in *Ectocarpus*, reflected by strong intra-chromosomal interactions and clear boundaries between chromosomes (see Figure 1B). We found a significant enrichment of inter-chromosomal interactions involving chromosomes 1, 12, 14, 20, and 27 (Figure 2A), suggesting a propensity for these chromosomes to establish stronger contacts compared to oth-ers. Furthermore, strong contacts among telomeric regions of different chromosomes, as well as contacts among centromeric regions (see below) were widespread on the Hi-C map (Figure 2B).

**Figure 2.**
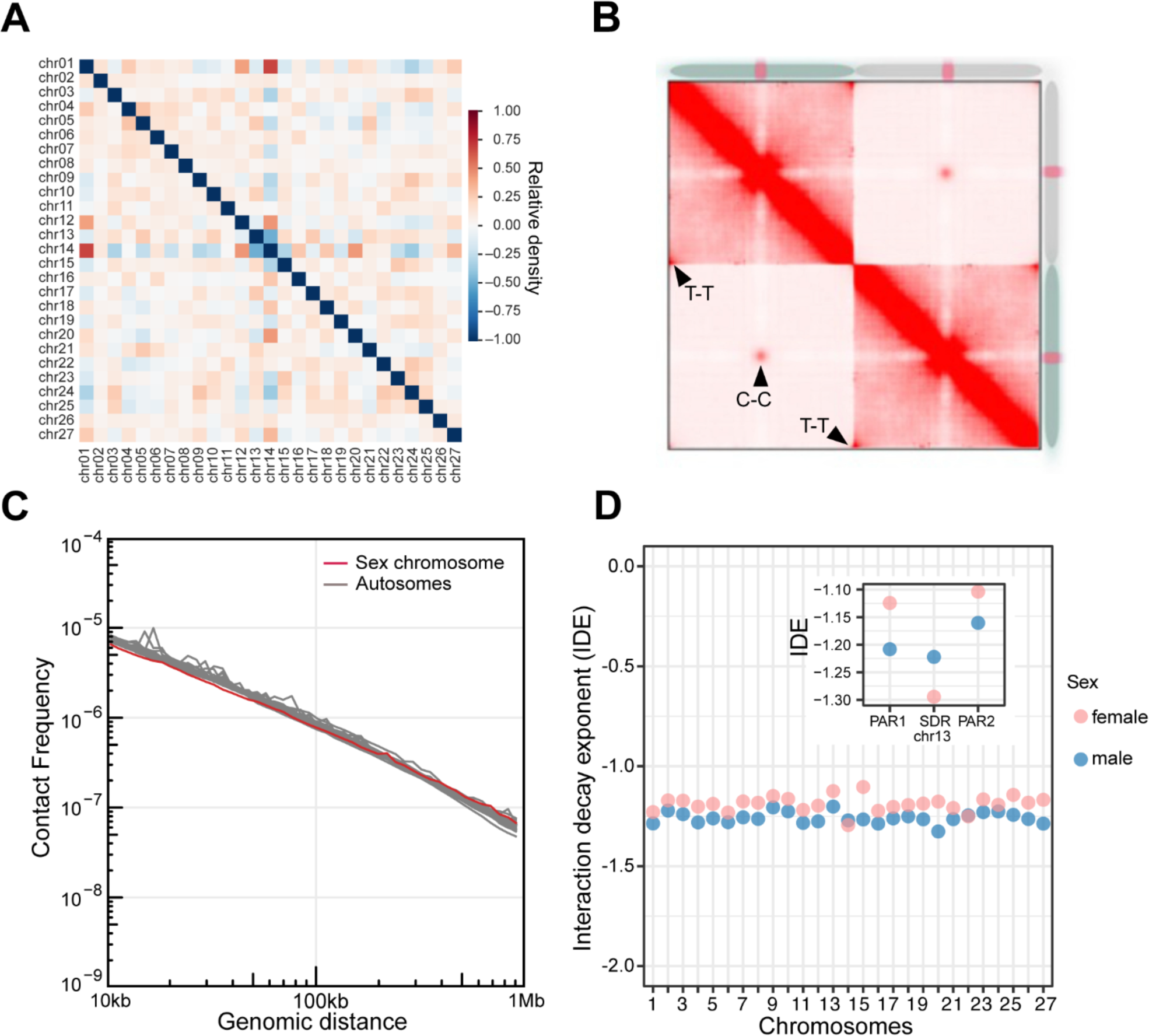
3D chromatin architecture of Ectocarpus revealed by Hi-C data. (A) Pair-wise averaged inter-chromosome con-tacts of Ectocarpus male at 10k resolution. (B) Analysis of aggregated intra- and inter-chromosomal contacts. T-T: telomere to telomere interactions; C-C: centromere to centromere interactions. (C) Global folding patterns of each of the male Ectocar-pus chromosomes reflected by contact frequency as a function of genomic distance (Ps). (D) IDEs of each autosome and sex chromosome regions in Ectocarpus male and female. Normalized Hi-C matrices at a resolution of 10 kb at distance range of 10 kb to 500kb were used to calculate IDEs. Figure 3: High-resolution contact probability map reveals the higher-order organ-ization of the Ectocarpus genome.

Next, we computed each chromosome’s chromatin contact probability as a function of genomic distance to ex-amine *Ectocarpus* chromosome packing patterns. As expected, we observed a decline in contact frequencies as genomic distances increased (Figure 2C). Next, Interaction Decay Exponents (IDEs), which describe how fast in-teraction frequencies drop with increasing physical genomic distance, were computed to characterize chromatin packaging ^34,35^. We found that for each of the *Ectocarpus* chromosome, interaction frequencies decayed in similar power-law functions with IDE values between 10kb and 500Kb (Figure 2D). However, the IDE values in SDRs and PARs of sex chromosomes showed noticeable variation, suggesting differences in local chromatin packing in these regions (see below).

One prominent feature of animal and plant genomes is the organization of chromatin into TADs, characterized by preferential contacts between loci inside the same TAD and strong insulation from loci in adjacent TADs ^20,36^. TADs regulate enhancer-promoter contacts and gene expression ^37^. Intriguingly, we did not observe conspicuous TADs patterns in any of the *Ectocarpus* chromosomes upon zooming into the Hi-C map (Figure S6). Note that *Ectocarpus* has a similar genome size to the land plant *Arabidopsis thaliana*, which also does not exhibit classical TAD struc-ture, but rather TAD-like domains that are moderately insulated from flanking chromatin regions ^25,38^ and is con-sidered as an outlier species concerning plant TAD formation ^39^.

### A/B compartment dynamics in males versus females

Spatially distinct nuclear compartments are a prominent feature of 3D chromatin organization in eukaryotes ^34^. A/B compartments, which generally correlate with active and repressed chromatin, respectively, can be identified by applying Principal Components Analysis (PCA) of the correlation heatmap yields the first eigenvector (EV1), (PCA, Lieberman-Aiden et al. 2009). We applied PCA to individual chromosome’s Hi-C maps normalized at 10kb bin size to identify the two spatial compartments (Figure 3A). The compartment that displayed stronger inter-chromosomal chromatin contacts was called ‘A’, whereas B compartment had lower inter-chromosomal contacts (Figure 3B). Interestingly, the genomic regions bearing the centromeres corresponded to the A compartment (Figure 3A, B). Further PCA analysis on the A compartment indicated that centromeres formed distinct sub-com-partments, which were spatially separated from the rest of the A compartment regions (Figure S7).

**Figure 3:**
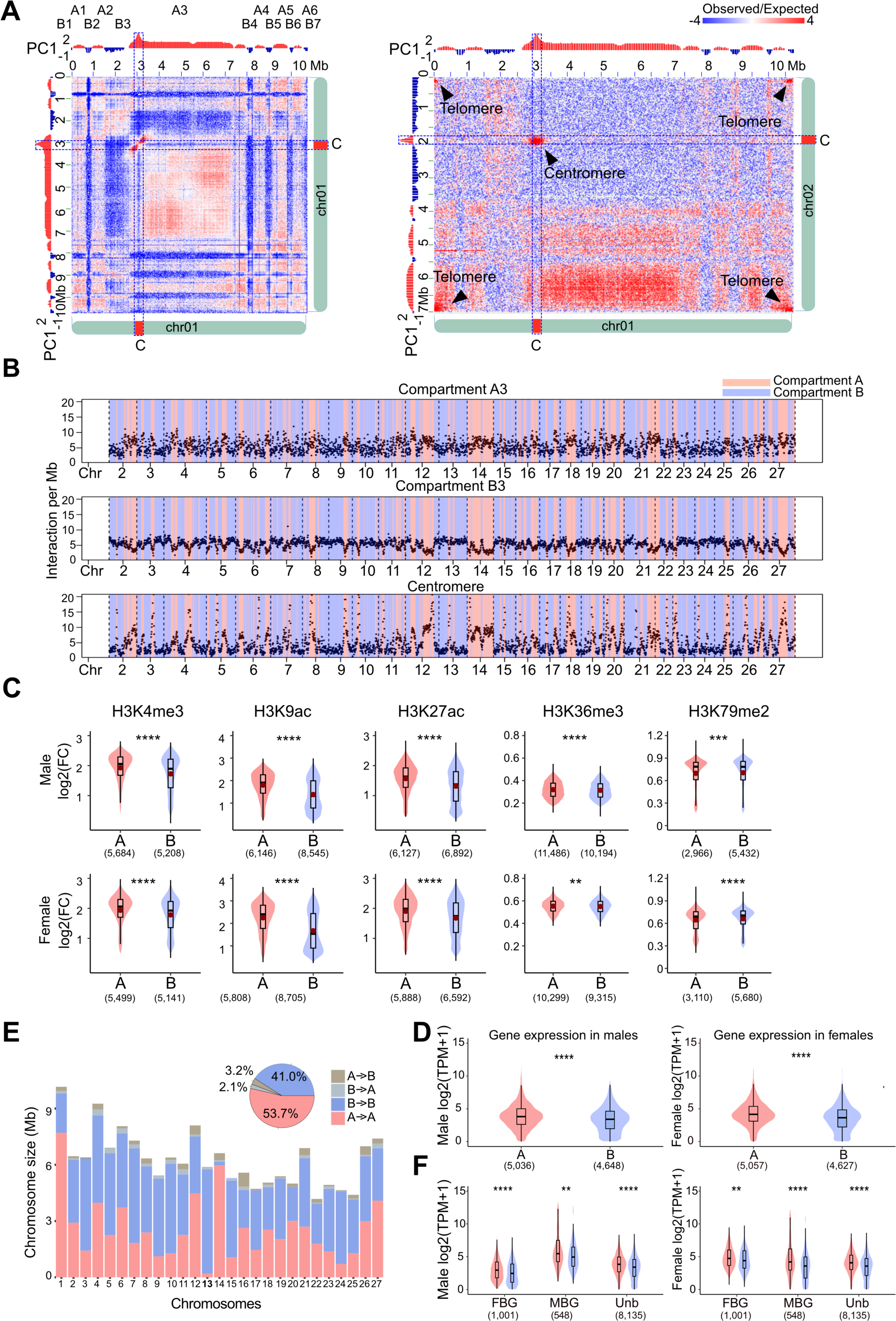
High-resolution contact probability map reveals the higher-order organization of the Ectocarpus genome. (A) Compartment A/B annotation based on principal component analysis. PC1 stands for the first principal component. The right panel shows inter-chromosomal contact patterns of A/B compartment regions between chromosomes 1 and 2. (B) Inter-chro-mosomal contacts of selected chromosome 1 regions with other chromosomes. The plots describe inter-chromosomal con-tacts belonging to the compartment A3 (top), B3 (middle), and centromere regions (bottom). The A/B compartment annota-tion of individual chromosomes is indicated with different colors. (C) Comparison of histone modifications, represented as fold changes of regions enriched with selected histone marks. For each histone mark, the enriched regions are grouped according to A/B Compartments of Ectocarpus male and female. Mean value of log2(fold change) is represented by a red dot in each boxplot. Wilcoxon test, NS: p > 0.05, *: p < 0.05, **: p < 0.01, ***: p < 0.001, ****: p < 0.0001. Numbers in brackets represent the number of peaks of the corresponding histone ChIP-seq data. (D) Levels of gene expression in compartment A and B in males and females. p-values of Wilcoxon test, **: p < 0.01, ***: p < 0.001, ****: p < 0.0001. Numbers in brackets represent the number of genes. (E) Sizes of conserved and switching A/B compartment regions in male and female Ectocarpus genomes. “X ->Y” indicates compartment annotation in male (“X”) and female (“Y”). The pie chart indicates pooled data from all chro-mosomes. (F): Expression of sex biased genes (SBG) in compartment A/B regions. MBG: male biased gene; FBG: female biased gene; Unb, unbiased gene. p-values of Wilcoxon test, **: p < 0.01, ***: p < 0.001, ****: p < 0.0001. Numbers in brackets represent the number of genes.

Although different chromosomes had different proportions of compartment A and B, we noticed that the U and V sex chromosomes (chromosome 13) exhibit large stretches of B compartment regions (Figure 3B), suggesting they have a distinct configuration compared to autosomes overall (see below).

The *Ectocarpus* genome has been reported to have various histone post-transcriptional modification (PTMs) associated with gene transcriptional activities ^40,41^. We therefore asked whether chromatin associated with different A/B compartments exhibited different histone modification profiles. To this end, we used published epigenomic datasets for a range of histone PTMs from the same strains (Ec560, Ec561) ^42^ and mapped the ChIP-seq datasets to our V5 genome. We found that for both male and female genomes, the histone PTMs associated with active gene expression, such as H3K4me3, H3K9ac, H3K27ac and, most conspicuously, H3K36me3, were significantly enriched in the A compartment (Figure 3C). Conversely, H3K79me2, a histone mark associated with repressed chromatin in *Ectocarpus* ^40,41^, was enriched in the B compartment (Figure 3C). Furthermore, genes located within A compartment regions exhibited higher expression levels than those in the B compartments (Figure 3D). These observations are in agreement with chromatin organization patterns reported in other multicellular eukaryotes where the A/B compartments are enriched with euchromatic and heterochromatic chromatin, respectively^23,34^.

The A/B compartment assignment of the male and female *Ectocarpus* genomes was highly similar; nonetheless, 5.3% of the *Ectocarpus* chromatin exhibited different A/B compartment identities in male and female Hi-C maps (Figure 3E). In animals, compartment status and boundaries may change during cell differentiation and correlate with changes in gene expression profiles ^43^. We therefore investigated whether such changes in compartment annotation were associated with the expression patterns of sex-biased genes (SBG), i.e., genes that show a significant change in expression in males versus females ^44,45^. We used RNA-seq datasets^42^ and identified 2,069 SBGs (see methods for details, Table S7). Depending on the expression preference, these SBGs were further annotated as male- and female-biased genes (MBGs and FBGs), respectively. Compared with the distribution of unbiased genes concerning A/B compartment annotation, SBG were not enriched in the regions where A/B compartment identity changed (Chi-square test p=0.0649), suggesting that the regulation of SBG expression is largely independent from the chromosomal conformation. MBG in males were upregulated when in compartment A compared to FBG, and the opposite was true in females (Figure 3F), and we noticed that whilst MBG in males still show greater expression associated to the A compartment, the overall expression levels were higher, regardless of the compartment. Therefore, the patterns of expression of SBG were correlated with their association to histone PTMs and to the specific 3D chromatin organization in males versus females.

### U and V sex chromosomes and autosomes adopt distinct conformations

The sex-specific high resolution genomic maps were then used to compare the sub-nuclear 3D genomic architec-ture of the U and V sex chromosomes. The U and V sex-specific regions (SDR) have been identified and character-ized previously^9,46,47^ but their largely repeat rich nature has prevented their full assembly. In the *Ectocarpus* V5, the V and U chromosome had a total length of 7.16 Mb and 7.23 Mb respectively (see Figure 1). In *Ectocarpus*, U and V are largely homomorphic with a small region that is non-recombining (SDR) and therefore largely divergent between male and female^9,47^) (Figure S8A). The male and female SDR of the *Ectocarpus* V5 genome feature no gaps. We also noticed that compared to the V2, the female SDR has increased in physical size (Figure S8B). This was mainly due to the addition of repeats in the new assembly (V2 had 34.7% of repeats and V5 68.3% of repeats in the U-SDR). The small SDRs are flanked by large pseudoautosomal regions (PARs), which recombine at meiosis ^9,48^. Structural analysis using our new assembly confirmed that U and V sex chromosomes display unique charac-teristics compared with autosomes, including lower GC content, higher repeat content and lower gene density ^9,48^ and a largely repressive chromatin landscape ^41^ (Table S4, Figure 4A). We then used the 2 kb resolution Hi-C map to investigate the 3D structure of the sex chromosomes in the *Ectocarpus* nucleus. Intriguingly, the U and V sex chromosomes exhibited a distinct 3D architecture compared to autosomes, with their central, sex-specific regions (SDRs) both being insulated from the flanking PAR regions (Figure 4B, see also Figure 2D), with high intra-chromosomal contacts in the 3D space. We also noticed that both U and V SDRs spanned the centromeres (Figure 4B).

**Figure 4:**
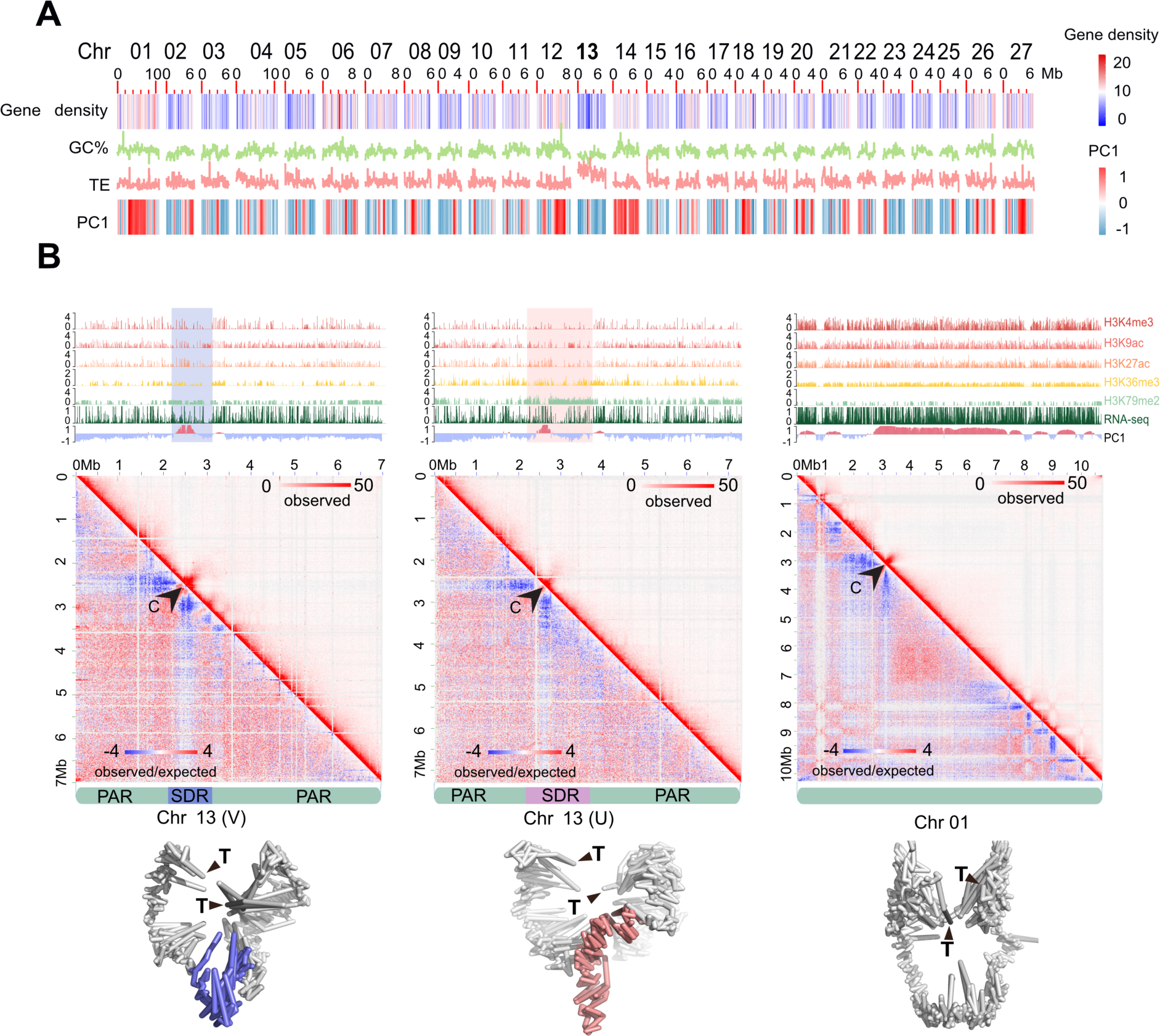
U and V sex chromosome 3D architecture. (A) Plot showing gene density, GC content, TE density in 100kb windows and compartment A/B (PC1) in 10kb windows across Ectocarpus chromosomes. Chromosome 13 is the sex chromosome. (B) Hi-C map and simulated 3D configurations of sex chromosomes at 10k resolution. SDRs in the simulated male and female chromosomes are colored in blue and red, respectively, and telomeres are labeled with black triangles. In each panel, the black arrowhead indicates centromere. The tracks above each Hi-C map show A/B compartment annotation (PC1), gene-expression (RNA-seq), and various histone modification ChIP-seq.

### *Ectocarpus* centromeres are distinguished by specific LTR retrotransposons

To determine the structure and precise locations of the *Ectocarpus* centromeres, we analysed the sequence characteristics of the chromosomal regions delineated by centromere-to-centromere interactions (see Figure 2B, Figure 3A, 3B). Regional centromeres vary extensively among eukaryotes, with common structures including short non-repetitive AT-rich regions, transposon-rich regions spanning tens to hundreds of kilobases, and megabase-scale satellite arrays (Talbert and Henikoff 2020). We first searched for any specific repeat families that were i) enriched in the putative centromeric regions, and ii) common to all chromosomes. This revealed two retrotransposon families that are almost exclusively restricted to a single highly localised, gene-poor, and repeat-rich region on each chromosome (Figure 5A, Figure S9). The most abundant of the two elements is a 6.6 kb *Metaviridae* (i.e. *Ty3*/*Gypsy*) long terminal repeat (LTR) retrotransposon, which encodes Gag and Pol on a single open reading frame of 1,699 aa and can be found as full-length copies flanked by 4 bp target site duplications (Figure 5B). The second is presumably a related LTR element, although it is only present in degraded fragments, and we were unable to recover an internal protein-coding region. The two retrotransposon families share a ∼64 bp region of homology that includes the polypurine tract that immediately precedes the 3’ terminal repeat (Figure 5B). We name these elements *ECR-1* and *ECR-2* for *<u>E</u>ctocarpus* <u>C</u>entromeric <u>R</u>etrotransposon.

**Figure 5.**
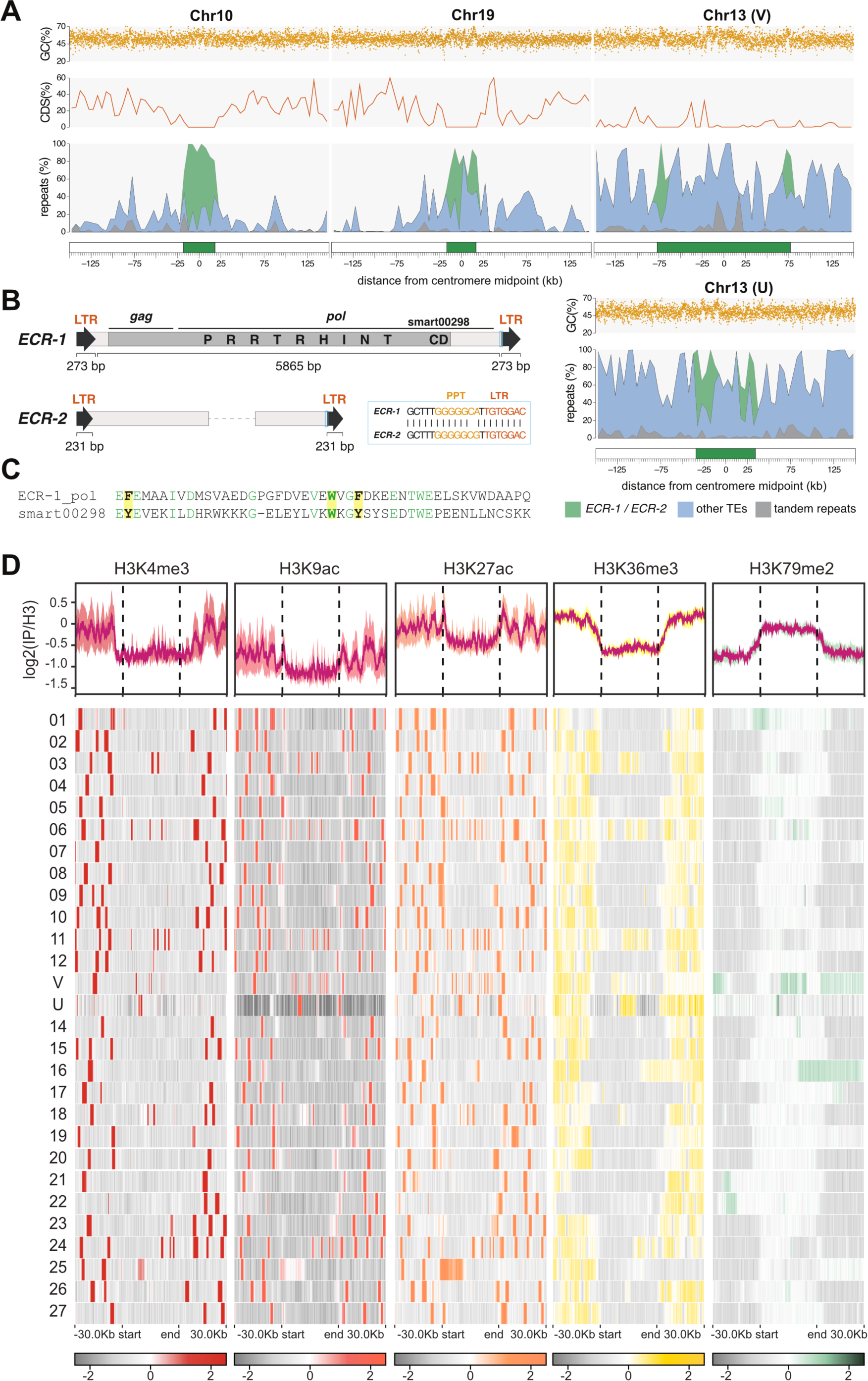
Ectocarpus centromeres and centromere-specific retrotransposons. (A) The centromeric regions of select chro-mosomes and the ECR retrotransposons. Putative centromeres and flanking regions for four chromosomes, including the U chromosome from the female genome assembly. The centromere (green box) is defined as the region from the first to the last copy of ECR elements. The repeats panel is shown as a stacked area plot, and the percentage of each repeat type is plotted in 5 kb windows. Coding sequence (CDS) density is plotted in 5 kb windows, and GC content is plotted in 100 bp windows. For all chromosomes see Supplemental Figure S9, and for genomic coordinates see Table S6. (B) Schematics of the ECR retrotrans-posons. The light blue boxes highlight the conserved region between ECR-1 and ECR-2, and a partial alignment of this region is shown (PPT = polypurine tract). Only 5’ and 3’ fragments of ECR-2 were recovered, and the dashed line represents protein-coding sequence that is presumably missing. The domains shown on the ECR-1 protein are: PR = protease, RT = reverse tran-scriptase, RH = RNaseH, INT = integrase, CD = chromodomain. (C) Alignment of ECR-1 chromodomain and SMART chromo-domain curated model (smart00298). Conserved amino acids are coloured green, and the three aromatic amino acids that are responsible for recognition of histone methylated-lysines are highlighted in yellow. (D) Histone mark signal (log2(IP/H3)) in the putative centromeres and the surrounding regions (30Kb). Plots were computed from male data and multi-mapping method with a bin size of 100bp. On top, profiles of histone marks around the centromeres. The dark pink line stands for the result and the thick light-colored line behind stands for the standard error. Note that the U chromosome was added to the male heatmap for simplicity. For the whole heatmap (separately done for male and female) see Supplemental Figure S10A.

Notably, the *ECR-1* polyprotein features a C-terminal chromodomain fused to the integrase domain (Figure 5B, C). Chromodomains recognise and bind histone methylated-lysines via a cage tertiary structure that is formed by three aromatic residues ^49^, all of which are conserved in the *ECR-1* polyprotein (Figure 5C).

Defining the putative centromeres as the region between the first and last *ECR* element, lengths range from only 6.8 kb on chromosome 25 (essentially a single copy of *ECR-1*) to 153.9 kb on the male chromosome 13 (i.e., chro-mosome V), with a median of 38.6 kb (Table S6). On average per centromere, 33% of bases are contributed by *ECR-1*, 7.3% by *ECR-2*, and 34% by other interspersed repeats that are not exclusive to these regions. Tandem repeats constitute only 3.5% of the putative centromeres, relative to 6.7% elsewhere in the genome. Although genes were generally absent from these regions, certain chromosomes feature a small number of genes distrib-uted among the *ECR* copies (e.g. chromosome V, Figure 5A). The GC content of the putative centromeres (52.7%) is only marginally lower than the rest of the genome (53.5%). However, this is partly driven by the GC content of *ECR-1* (58.2%), and several chromosomes do feature short AT-rich sequences within the putative centromeres (e.g., chromosome 19, Figure 5A). As expected, following their evolutionary independence, the putative centro-mere of the female U chromosome differs substantially in length and composition relative to the V chromosome.

To further characterize the *Ectocarpus* putative centromeres, we analyzed the associated chromatin pattern using ChIP-seq data (Figure 5D, Figure S10A). Despite their assignment to compartment A, the putative centromeres exhibit a slight enrichment with the H3K79me2 mark. Furthermore, the surrounding regions of these putative centromeres contain prominent peaks of histone marks associated with active genes (H3K4me3, H3K9ac, H3K27ac and H3K36me3), consistent with their compartment assignment and the presence of flanking genes. Interestingly, on a few chromosomes, the H3K79me2 pattern extends beyond the boundaries of the ECR. This observation holds true when using a different mapping method (removing multi-mapping reads) for chromo-somes 1, 16, U, and V (Figure S10B). These regions were highly enriched with transposable elements (TEs). Nota-bly, the core of the V centromere exhibited a strong signal in H3K79me2 marks, which is not retrieved in the U chromosome, except one strong spike of H3K79me2.

### An inserted (endogenous) viral element exhibits a unique chromatin conformation

Marine filamentous brown algae of the order Ectocarpales frequently carry endogenous giant viruses with large double-stranded DNA genomes ^50^. *Ectocarpus sp.7*, in particular, has been shown to harbor such type of endoge-nous viral element inserted in chromosome 6, derived from the *Ectocarpus* phaeovirus EsV-1^29,51,52^. We confirmed the presence of an endogenous viral element (that we name Ec32EVE) localizing to chromosome 6 in our V5 *Ectocarpus* genome (Figure 6A, see also Figure 1A,). The Ec32EVE is 399 kbp long and contains 199 genes, and is covered with a large domain of the repression-associated mark H3K79me2 previously shown to be associated with the silencing of transposable elements in *Ectocarpus*^40,41^. The Ec32EVE region exhibits a depletion of activa-tion-associated histone marks H3K4me3, H3K9ac, H3K27ac and H3K36me3 (Figure 6A). Consistent with this het-erochromatic landscape, RNAseq analysis showed negligible expression throughout the entire Ec32EVE region (Figure 6A), highlighting the silent nature of the potentially coding regions within the endogenous viral element. The chromosome 6 Hi-C map further revealed high levels of compaction and insulation, associated with the viral insertion region (Figure 6A). Remarkably, the Ec32EVE region displayed strong long-range contact with telomeres in the nuclear 3D space (Figure 6B). We asked whether this observation was related to the highly heterochromatic nature of the Ec32EVE region. However, other *Ectocarpus* genomic regions equally marked with long stretches of H3K79me2 did not necessarily cluster in 3D with telomeres (Figure S11). It appears, therefore, that the Ec32EVE insertion, rather than the epigenetic footprint of this region *per se*, is implicated in the unique 3D structure of this region.

**Figure 6.**
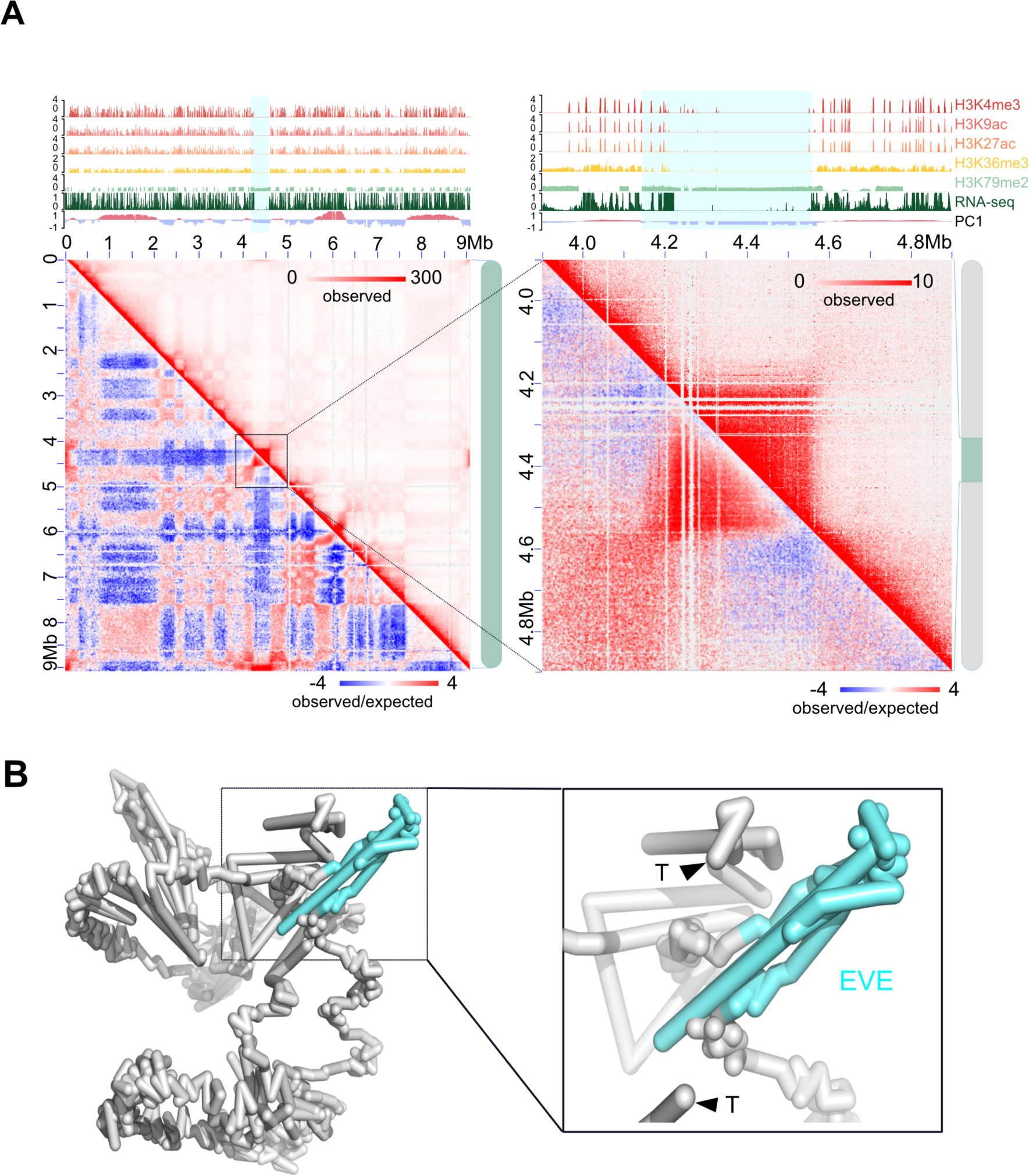
Virus insertion region (EVE) is insulated and shows strong interactions with telomeres. (A) Hi-C map of chromosome 6. The zoomed-in region to the right contains the EVE (4.2-4.6 Mb). The tracks above each Hi-C map show A/B compartment annotation (PC1), gene-expression (RNA-seq), and histone PTMs ChIP-seq tracks. A blue shade marks the region of the Ec32EVE. (B) Simulated 3D configurations of chromosome 6 at 10k resolution. The EVE region is colored in aquamarine, and the telomeres are labeled with black triangles. To the right, the Ec32EVE region is zoomed-in to highlight the long-range contacts with the telomeres.

## Discussion

High-quality and complete reference genome assemblies are fundamental for the application of genomics to a range of disciplines in biology, from evolutionary genomics, genetics to biodiversity conservation. Here, we obtained a highly accurate and nearly complete assembly of the reference genome of the brown alga *Ectocarpus*, a model organism for this key group of eukaryotes. The *Ectocarpus* V5 assembly includes telomeres for most chro-mosomes, very few gaps, and therefore provides a new reference genome for the scientific community.

Chromosome folding patterns vary across lineages^53^. For example, in many plant species with relatively large genomes, chromosomes adopt Rabl configuration during interphase, in which centromere or telomere bundles are associated with opposite faces of the nuclear envelope. For chromosomes with Rabl configuration, their Hi-C maps display a characteristic belt that is perpendicular to the primary diagonal. *Arabidopsis*, in contrast, presents a Rosette configuration ^54^, where the Hi-C maps feature conspicuous long-range intra-chromosomal contacts due to the formation of megabase-size loops. None of these features were found in the *Ectocarpus* Hi-C map, suggest-ing that its chromatin adopts a non-Rabl and non-Rosette configuration. Chromatin arrangement of interphase chromosomes in *Ectocarpus* involved telomeres of all chromosomes and centromeres of all chromosomes clus-tering together. Therefore, despite different linear genome architectures and centromere sequence composi-tions, centromere interactions appear to be a pervasive feature in eukaryotes, from plants and animals to brown algae.

TADs, whose boundaries partition the genome into distinct regulatory territories, are a prevalent structural fea-ture of genome packing in animal and plant species, but our observations showed that TADs are not prominent in the *Ectocarpus* genome. Note that *Ectocarpus* has a relatively simple morphology with a reduced number of cell types. The Hi-C maps, thus, are likely to faithfully represent the interphase chromatin structure of male and female *Ectocarpus* rather than an average conformation across multi-cell types as in other more complex organ-isms. This feature allows us to conclude that *Ectocarpus* has a non-Rabl chromatin conformation and does not exhibit TADs at a local level. The *A*. *thaliana* genome is another example in which TADs are absent^25^ and this feature is thought to be related to *A*. *thaliana* genome small size, high gene density, and short intergenic regions. Given that *Ectocarpus* also has similar genomic characteristics, the absence of TADs in *Ectocarpus* supports the hypothesis that TADs may form when the genome size is above a certain threshold ^55,56^. Note that in *A*. *thaliana*, despite its genome not having clear TADs, over 1000 TAD-boundary-like and insulator-like sequences were found from Hi-C maps normalized with 2 kb genomic bins ^25^. These regions possess similar properties to those of animal TAD borders/insulators, i.e., chromatin contacts crossing insulator-like regions are restricted, and they are en-riched for open chromatin. The *Ectocarpus* genome, in contrast, is mainly partitioned in H3K79me2 rich and H3K79me2 poor regions, that largely define A and B compartments, but we did not find any evidence for canonical insulators nor ‘TAD-boundery-like’ regions. Note that CTCF is absent in the genome of *Ectocarpus,* similarly to yeast, *C. elegans* and plants ^57^.

The high resolution, sex-specific Hi-C maps of haploid individuals allowed us to examine the 3D structure of the U and V sex chromosomes in the interphase nucleus of *Ectocarpus* males and females. The U and V chromosomes are largely homomorphic, each containing a small, non-recombining region^9,47^ that harbors several dozen genes, including the master male-determining factor MIN^58^, and a largely heterochromatic landscape ^41^. Our *Ectocarpus* V5 yielded gapless SDRs and demonstrated that the U and V SDRs span the centromere. Linkage between mating type (MT) locus and centromeres is a common feature of haploid mating-type (MT) chromosomes in fungi. For example, in the *Microbotryum* fungi, recombination suppression links the MT determining loci to centromeres ^59,60^ and this is thought to help preserving heterozygosity and/or be beneficial under auto-fecundation, as it in-creases the degree of compatibility between gametes from the same individual. A similar process is unlikely to be operating in *Ectocarpus* because there is no intra-tetrad direct crossing; haploid spores disperse after meiosis, develop into male and female gametophytes and produce gametes at a later stage ^61^. It is therefore more conceivable that SDR linkage to centromere in *Ectocarpus* occurred due to expansion of the non-recombining SDR, during which the centromere was subsumed in this region likely via a large scale inversion ^47^.

What is the potential role of the sex chromosome 3D chromatin configuration? Among numerous steps required for gene expression, the spatial organization of the genome is known to modulate DNA accessibility to the transcriptional machinery and to promote contacts between genes and distant regulatory DNA elements such as enhancers^62^. In the case of *Ectocarpus*, the correct spatial and temporal window of transcriptional activation of genes contained within the SDR is critical to ensure sex determination and differentiation in the brown algal tissues. It is therefore likely that the tight transcriptional regulation of the SDR is achieved both by 3D chromatin remodeling in conjunction with histone PTMs and sRNA ^42^. Whilst the 3D chromatin configuration of animal and plants sex chromosomes remains largely elusive, it is well known that chromatin 3D structure is involved in repression of the silent MT loci in yeast *Saccharomyces cerevisiae* during mating type switching ^63^ ^64^. Therefore, it appears that modulation of mating type or sex chromosome architecture may play a significant role in controlling sex-specific features across eukaryotic lineages.

The *Ectocarpus* V5 assembly and high-resolution Hi-C map allowed us to examine centromeric sequences in this organism. We observed 27 unique centromere sequences occurring once per chromosome, a finding that helps to resolve nuclear genome organization and indicates monocentric regional centromeres. The centromeres of *Ectocarpus* may be categorised as transposon-rich and primarily composed of centromere-specific retrotransposons. However, although *ECR-1* presumably targets centromeric DNA (and *ECR-2* may have done so in the past), it remains to be determined whether the *ECR* elements constitute the epigenetic centromere. The putative centromeres are short relative to the transposon-rich centromeres of many other species ^65^, and we cannot rule out an association between cenH3 and short AT-rich sequences, as in diatoms ^66^. ChIP-sequencing of cenH3 will be required to distinguish between these possibilities.

The fusion of an LTR integrase to a C-terminal chromodomain is most widely known from the evolutionary ancient chromovirus clade of *Metaviridae* LTRs, where the presence of the chromodomain enables recognition of specific histone modifications and targeted insertion at associated genomic sites^67^. In plants, the CRM subclade of chromovirus LTRs contains many centromere-targeting families that accompany satellite arrays and constitute a major component of centromeric DNA^68^. Independent lineages of chromodomain-containing LTRs have been reported in Stramenopiles, including the Chronos *Metaviridae* elements of oomycetes^69^ and the *CoDi*-like *Pseudoviridae* (i.e. *Ty1*/*Copia*) elements of diatoms^70^. Interestingly, *ECR-1* does not appear to be a member of either the chromovirus or Chronos clades, and instead is most closely related to chromodomain-containing oomycete LTRs that are yet to be phylogenetically classified (e.g. *Gypsy-20_PR* from *Phytophthora ramorum*). We hypothesise that the chromodomain of *ECR-1* may enable centromere-targeted integration in *Ectocarpus*, either by recognition of cenH3 or another centromere-associated epigenetic context, implying evolutionary convergence with the CRM elements of plants. However, it is unlikely that the targeting mechanism is itself convergent, since the centromere-targeting CRM elements feature derived chromodomains that lack the three conserved aromatic amino acids ^68^, which are present in *ECR-1*.

Viruses that transcribe their DNA within the nucleus have to adapt to the molecular mechanisms that govern transcriptional regulation. The interaction between chromatin and viral directed modulation of chromatin are critical component of the viral-host interaction ^71^. However, the complexity of the higher-order organization of the host genome and its potential influence in the regulation of gene expression raises questions regarding the spatial arrangement of integrated viral DNA in the host’s genome. Phaeoviruses are latent giant double-stranded DNA viruses that insert their genomes into those of their brown algal (Phaeophyceae) hosts^50,52^. Remarkably, alt-hough about 50% of individuals in *Ectocarpus* field populations show symptoms of giant viral infection ^72^, the *Ectocarpus* strain used in this study has never been observed to produce virus particles, and Ec32EVE genes are transcriptionally silent ^29^. Here, we showed that the silencing of Ec32EVE genes correlated with deposition of large domains of repressive-associated chromatin mark H3K79me2, concomitant with depletion of activation-associ-ated marks. Moreover, the inserted giant viral element was associated to the B compartment, and adopted a highly insulated conformation in the 3D nuclear space, exhibiting strong long-range contacts with the telomeres. Whilst the detailed mechanisms underlying the relationship between giant virus latency and gene-silencing mech-anisms, including 3D architecture of the chromatin, remain to be determined, our study provides strong evidence for an interplay between 3D chromatin architecture, H3K79me2 domains and EVE gene silencing, as open new avenues to gain insights regarding the functional significance of these interactions.

## Supplemental Figures

**Figure S1.**
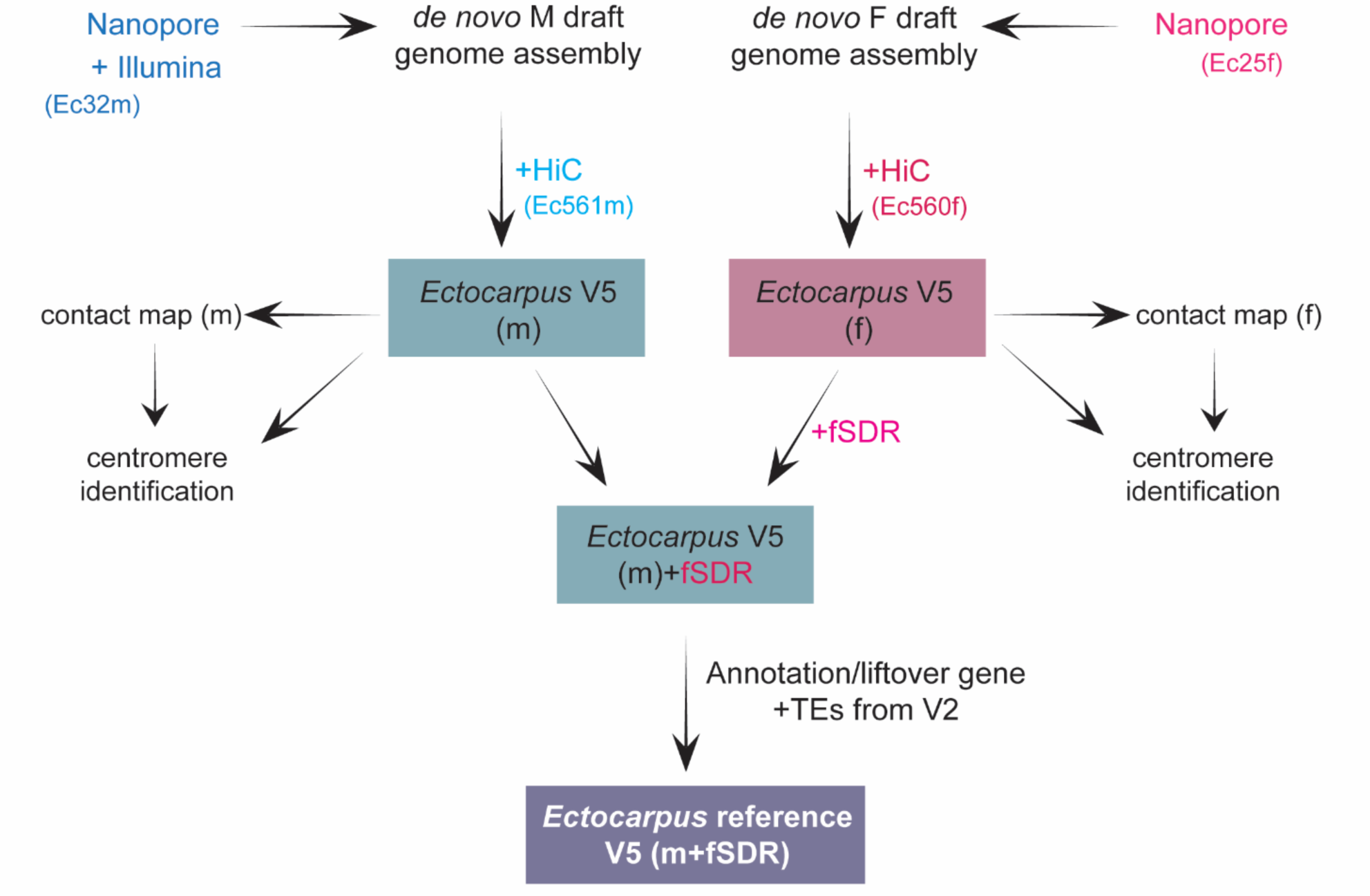
Schematic view of the approach used to reach a high-quality genome assembly of male and female Ectocarpus. De novo draft genomes from male and female siblings (Ec32 male and Ec25 female) were generated using Nanopore, and the male genome was polished using illumine reads. Genomes were further assembled using Hi-C data from Ec560 and Ec561 (near isogenic male and female lines ^41^. This resulted in the generation of male and female V5 genomes that were used for producing high resolution male and female contact maps. In order to have only one ‘reference’ genome, we chose to use the male V5 reference genome and complemented it with the female-specific sex-determining contig (SDR). Therefore, the final ‘reference’ Ectocarpus genome V5 is composed of high-quality male genome that includes both the male and the female SDR. Annotation of this V5 reference genome was performed by lifting over gene and TEs annotations from the Ectocarpus V2 genome ^28^. Centromeres were identified based on the contact maps and their annotation further refined (see methods for details). Ectocarpus strain numbers are given inside brackets (see also Table S1).

**Figure S2:**
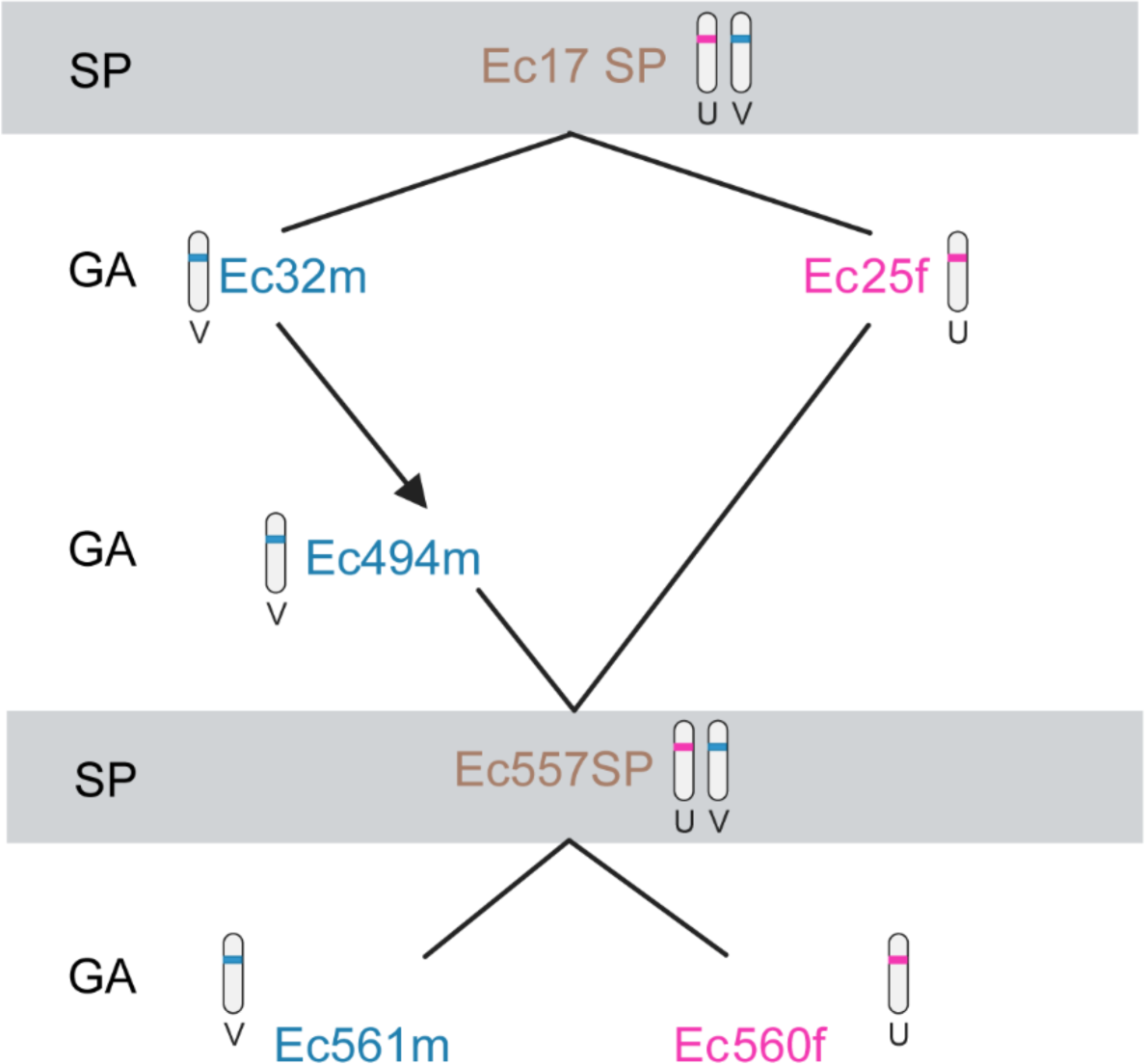
Pedigree of the Ectocarpus strains used in this study. SP, diploid sporophyte; GA, gametophyte; m, male game-tophyte; f, female gametophyte.

**Figure S3.**
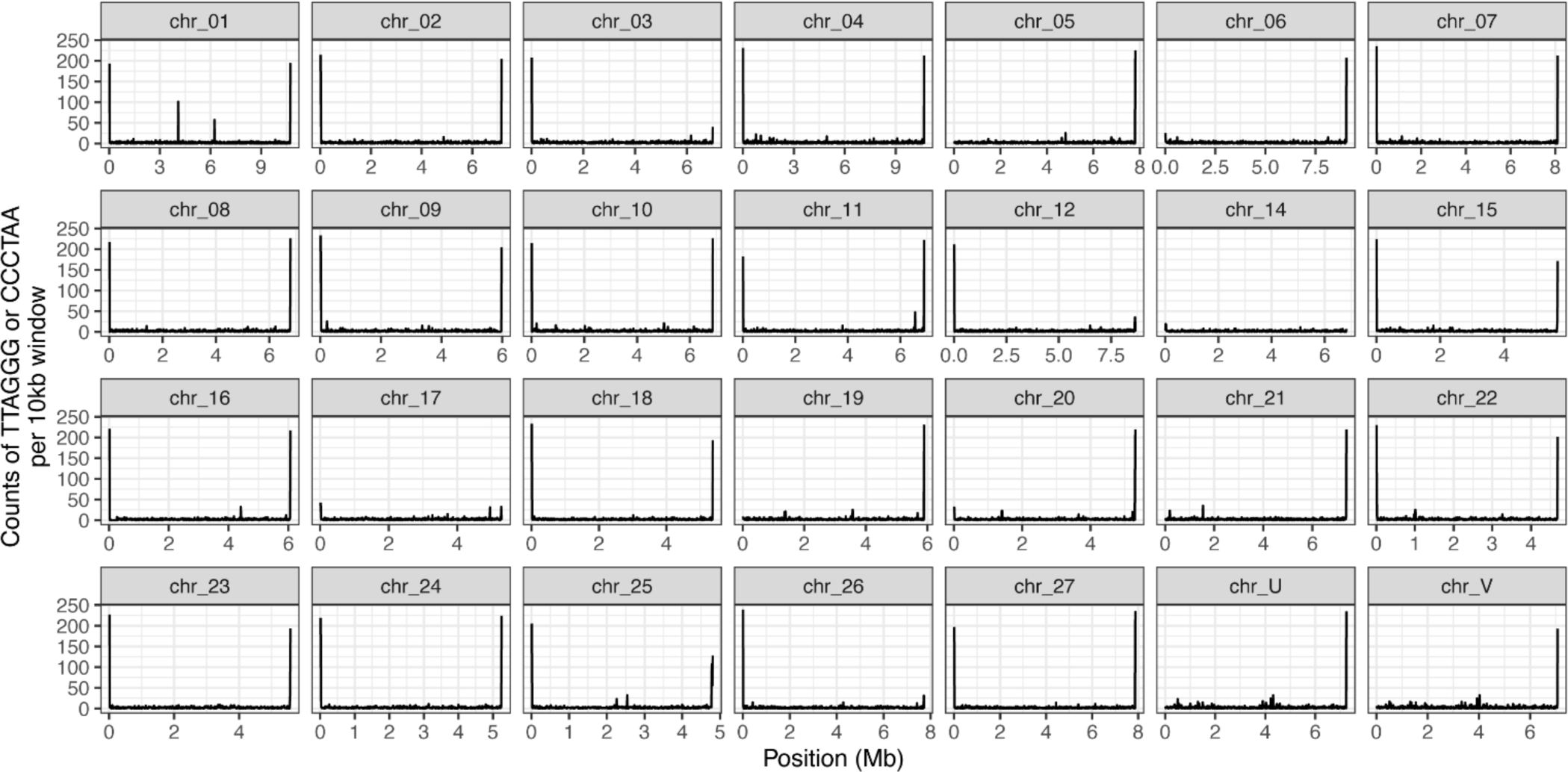
Distribution of telomere repeat motif (TTAGGG or CCCTAA) in haploid genome.

**Figure S4.**
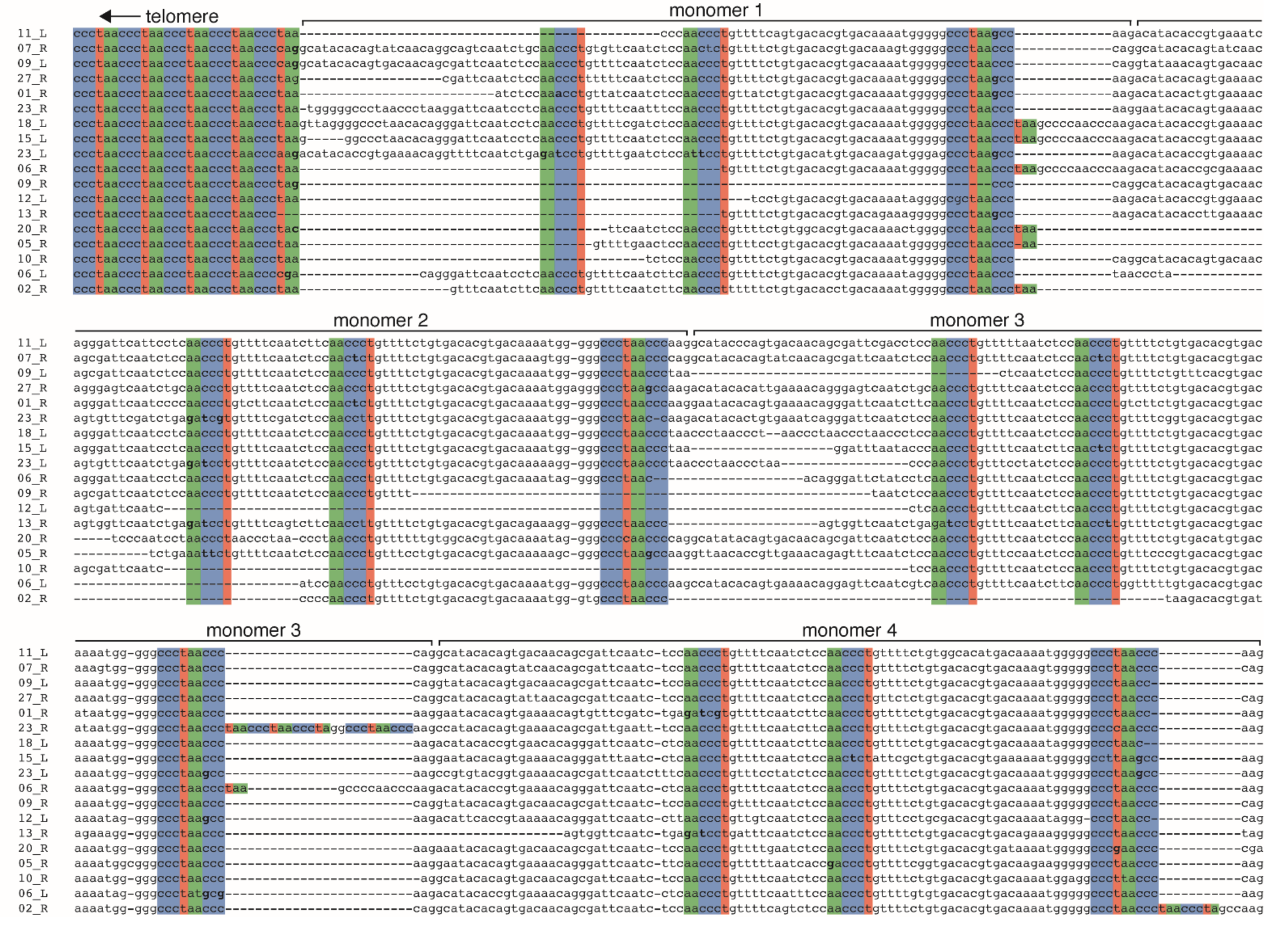
The organization of Ectocarpus subtelomeres. Alignment of 18 subtelomeres (e.g. 11_L is the left extremity of chromosome 11) showing the transition from the telomere to the subtelomeric satellite. The first four monomers of the ∼98 bp satellite are shown. Telomeres and telomeric motifs present in the subtelomeric satellite are colored.

**Figure S5.**
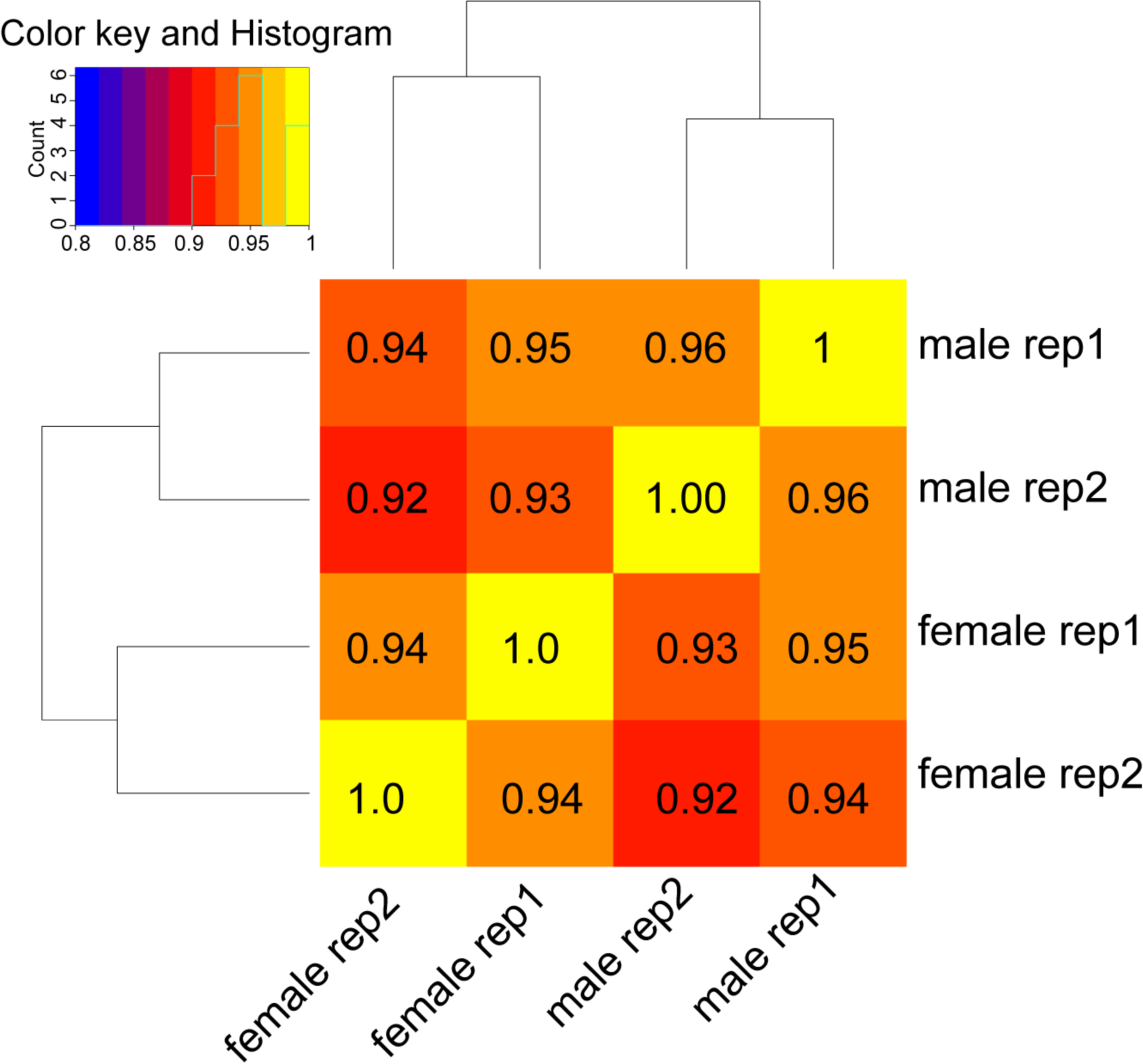
Quality control of biological replicates of Hi-C data. Pearson correction of biological replicates of Ectocarpus male and female Hi-C data at 10k bin size.

**Figure S6.**
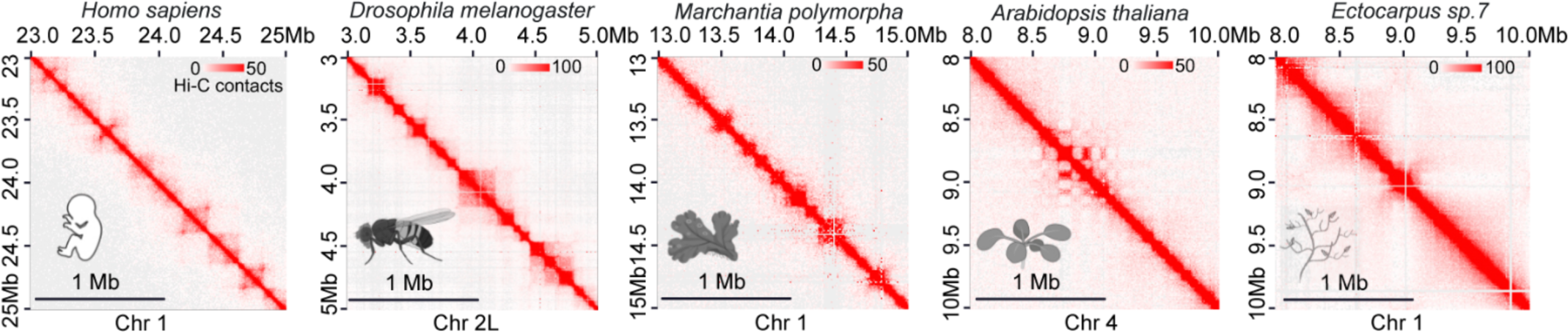
No prominent TAD patterns are observed in Ectocarpus. Examples of Hi-C maps from different species represent-ing TADs patterns in H. sapiens (chromosome 1,) Drosophila (chromosome 2L, ^73^), Marchantia (chromosome 1,^74^), Arabidopsis (chromosome 4,^38^) and Ectocarpus (chromosome 4, our paper). HiC maps of the different species were obtained using Juic-erbox at 10kb resolution. Note that TADs are not an obvious feature of Arabidopsis nor Ectocarpus genomes.

**Figure S7.**
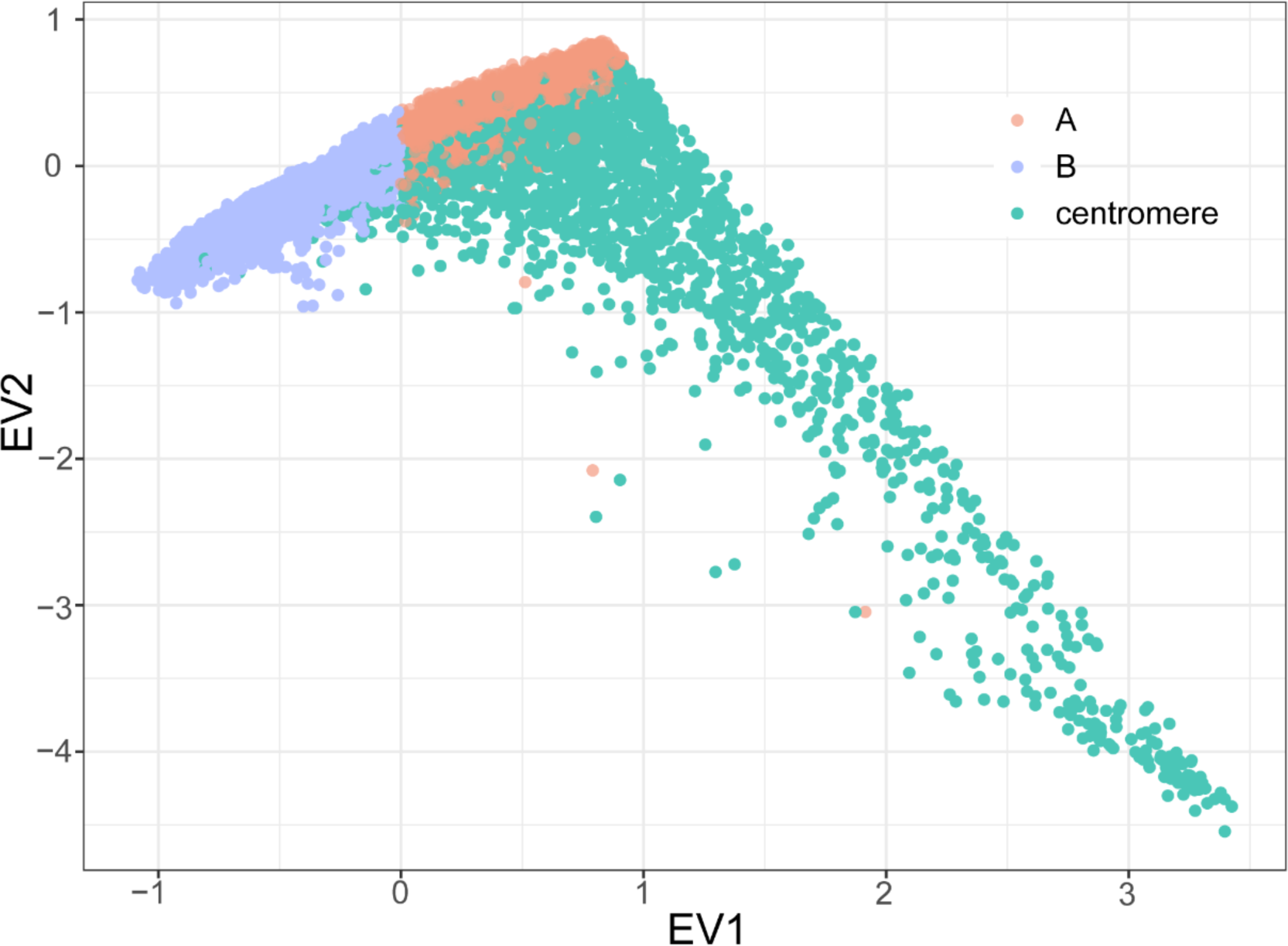
Centromeres form distinct sub-compartments. Centromere is represented as a sub-compartment, which could be separated from compartments by E1(PC1) and E2(PC2)

**Figure S8.**
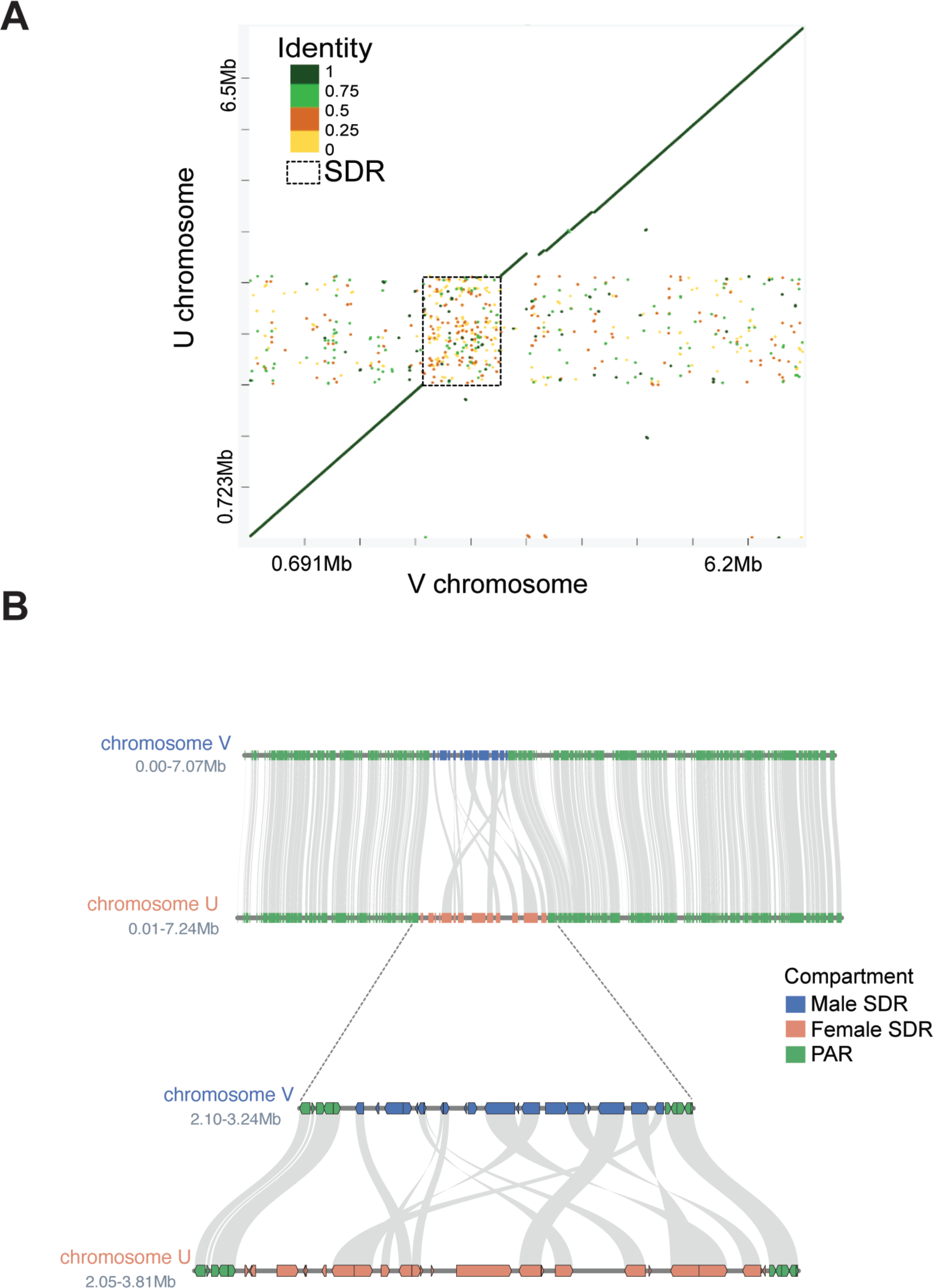
Sequence alignment of U and V sex chromosome. (A)The matches are presented as colored lines. The colors correspond to identity values that have been clustered in four groups (below 25%, between 25% and 50%, between 50% and 75% and over 75%), the dash box shows SDRs. (B) Synteny plot male vs female sex chromosome, including the newly, fully continuous male and female SDRs.

**Figure S9.**
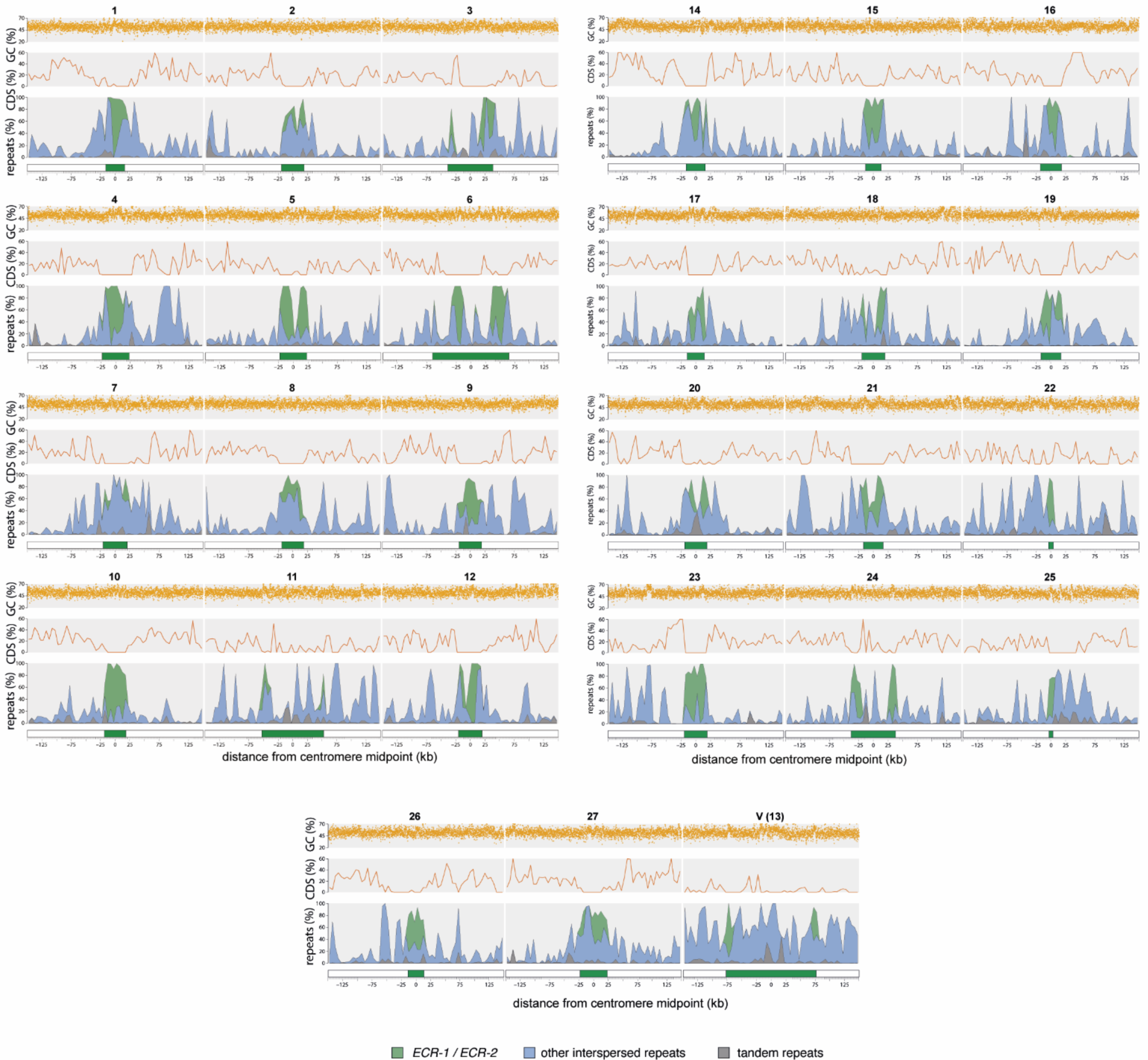
Sequence characteristics of Ectocarpus centromeres. Putative centromeres and flanking regions for all chromo-somes from the male Ec32 v5 assembly. The centromere (green box) is defined as the region from the first to the last copy of ECR elements. The repeats panel is shown as a stacked area plot, and the percentage of each repeat type is plotted in 5 kb windows. Coding sequence (CDS) density is plotted in 5 kb windows, and GC content is plotted in 100 bp windows.

**Figure S10.**
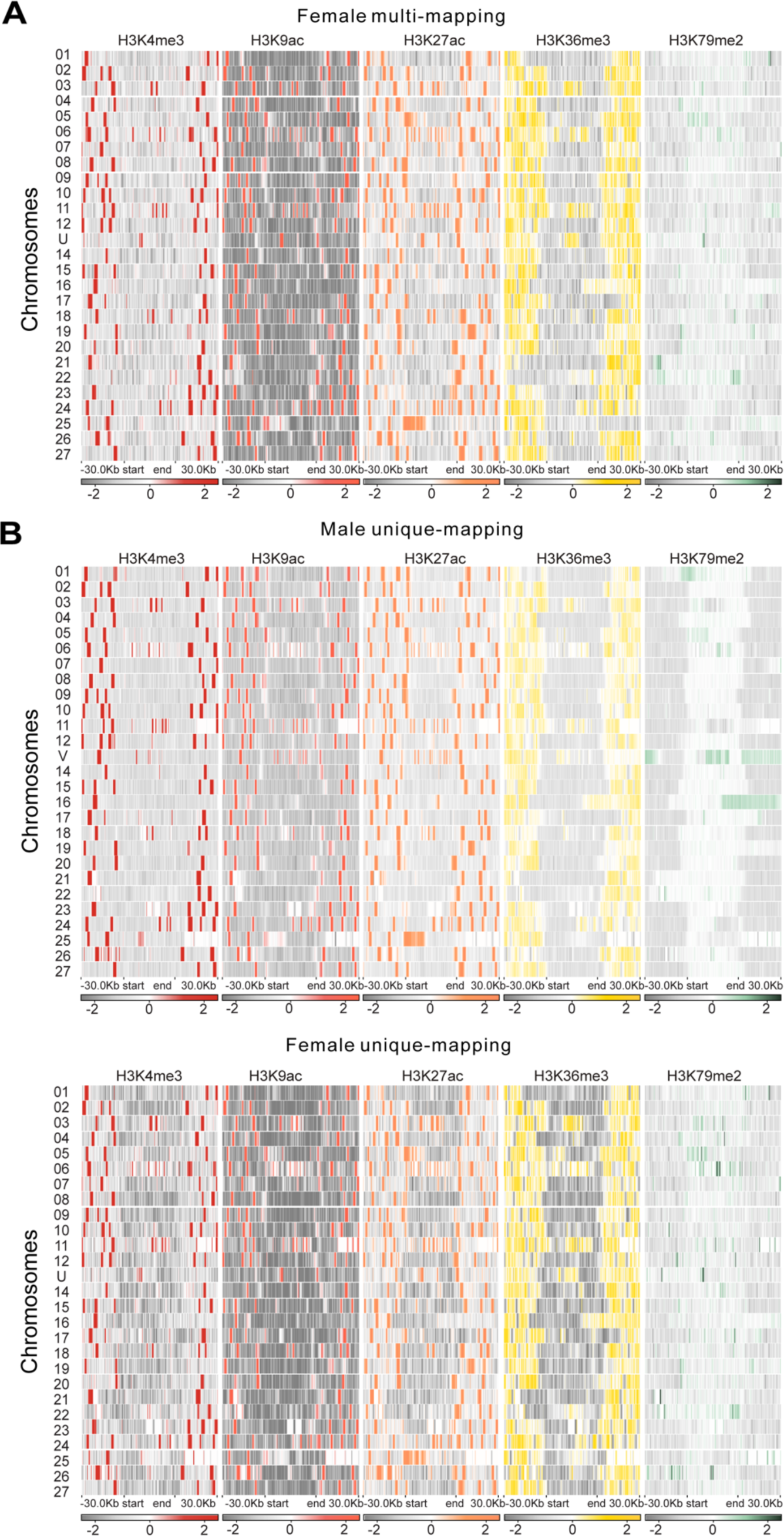
Heatmaps of histone marks around centromeres. For each heatmap, log2(IP/H3) is calculated on the putative centromere regions and 30Kb surroundings with a bin size of 100bp. (A) Heatmap from female data and multi-mapping method. (B) Heatmap from male and female data with unique-mapping method.

**Figure S11.**
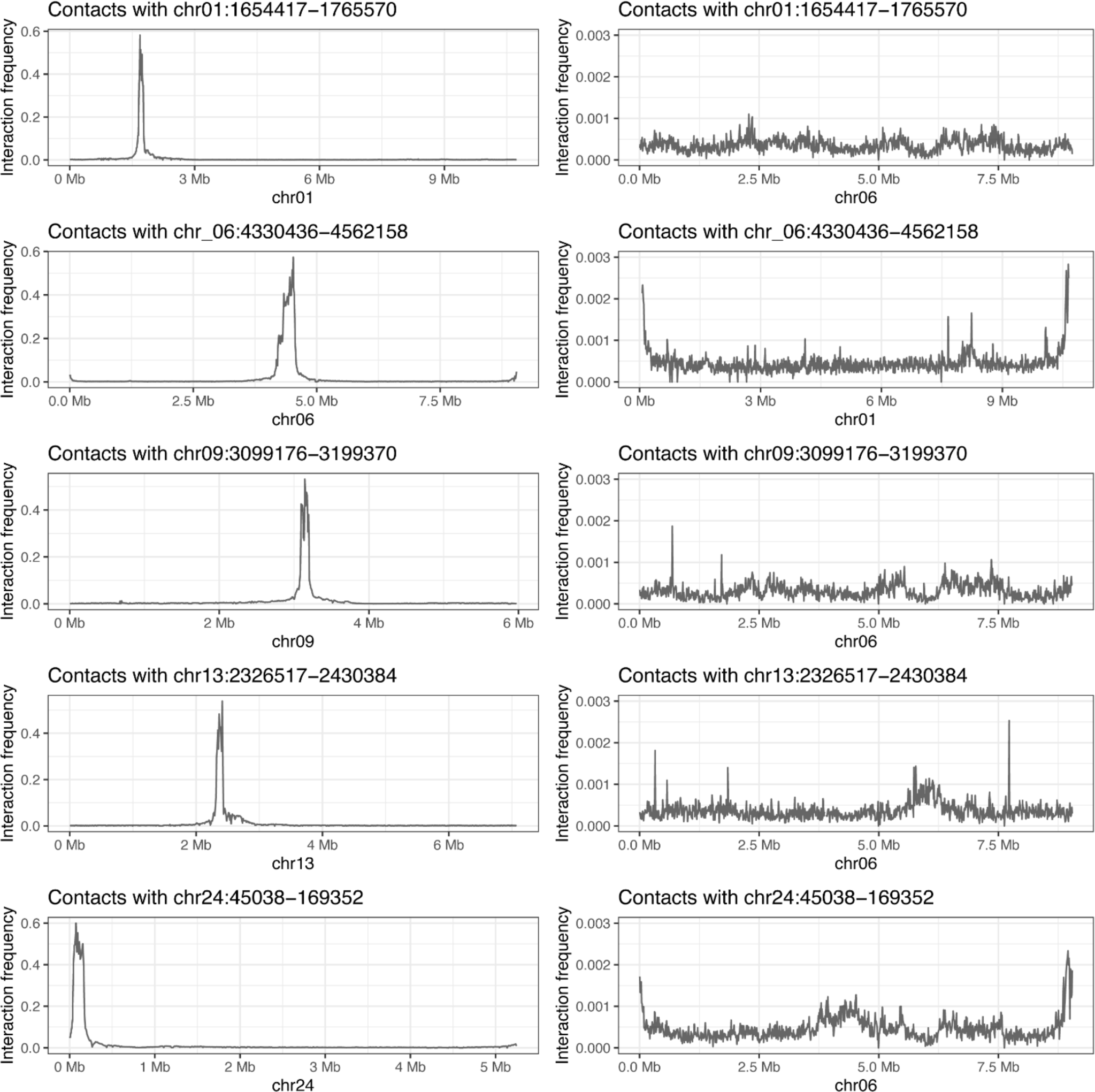
Examples of virtual 4C-like plot of H3K79me2 domains larger than 100 kb. Intra/inter chromosomal interaction frequency of H3Km79me2 domains.

## Supplemental Tables

Table S1. *Ectocarpus* strains used in this study. Table S2. Nanopore long reads statistics.

Table S3. Statistics of number of Hi-C reads in the different samples and replicates

Table S4. Genome statistics (chromosome level assembly). Note that Chromosome (chr) 13 is the sex chromo-some. Statistics are shown separately for male (V) and female (U) sex chromosomes.

Table S5. Statistics of complete, single, duplicated, fragmented, and missing genes computed by BUSCO v5.7.0

Table S6. Genomic coordinates and repeat densities for *Ectocarpus* putative centromeres. Centromeric coordi-nates were defined as the first to the last copy of ECR elements.

Table S7. Gene expression levels (transcripts per million, TPM) in the *Ectocarpus* male versus female samples used in this study (gametophytes)

Table S8. Number of sex biased genes (SBG) in compartments A and B in males and females.

Table S9: Comparative statistics of *Ectocarpus* sex determination region (SDR) annotations between the *Ectocar-pus* V2 and V5 assembly.

## Methods

### Brown algae culture

Algae were cultured as previously described ^75^. Briefly, *Ectocarpus* strains Ec32, Ec25, Ec561, and Ec560 were grown in autoclaved natural seawater (NSW) with PES at 14 °C with the light intensity of 20 μmol photons m^−2^ s^−1^ (12h light/12h dark). The medium was changed every week. Before collection, algae were treated with antibiotics: Streptomycin (25mg/L), Chloramphenicol (5mg/L), and PenicillinG (100mg/L) for three days to limit bacterial growth.

### Hi-C

An *in situ* Hi-C protocol of plants ^76^ was optimized for brown algae. *Ectocarpus* cultures were collected using a 40µm filter and fixed in 2% (vol/vol) formaldehyde for 30 min at room temperature, and the cross-linking reaction was quenched with 400 mM glycine. Approximately 50 mg fixed algae suspended in 1 ml nuclei isolation buffer (0.1% triton X-100, 125 mM sorbitol, 20 mM potassium citrate, 30 mM MgCl2, 5 mM EDTA, 5 mM 2-mercaptoeth-anol, 55 mM HEPES at pH 7.5) with 1X protease-inhibitor in a 2 ml VK05 tube, then homogenized by Precellys Evolution beads homogenizer (Bertin technologies) with the following settings: 7800 rpm, 30s each time, 20s pause each grinding cycle, repeat 5 times. Over 1 million nuclei were isolated and digested overnight by Dpn II, DNA ends were labeled with biotin-14-dCTP, then ligated by T4 DNA ligase enzyme. The purified Hi-C DNA was sheared by covaries E220 evolution and libraries were prepared using the NEBNext Ultra II DNA Library Prep Kit (NEB, no. E7645), and the average size of the library was detected by bioanalyzer, the final library was sequenced with 150bp paired-end reads on an Illumina HiSeq 3000 platform. Two biological replicates were performed for each strain.

### Nanopore sequencing

High molecule weight (HMW) DNA of *Ectocarpus* male (Ec32) and female (Ec25) were isolated using *OmniPrep*™ kit (G-Biosciences) with slight modifications. 500 mg of fresh collected tissue was dried and resuspended in 1ml lysis buffer, then homogenized using a Precellys mixer. Samples were incubated at 60°C for 1h with proteinase K, inversed every 15 min. HMW-gDNA was dried and eluted by 10mM ph 8.0 Tris-HCl, and incubated at 55°C for 30 min with 0.5 μL 10mg/ml RNaseA. The concentration of HMW-gDNA was quantified using an Invitrogen Qubit 4 Fluorometer, and molecule size distributions were estimated using a FEMTO Pulse system (Agilent). The sample was further cleaned and concentrated using AMPure XP SPRI paramagnetic beads (Beckman Coulter) at a DNA: bead volume ratio of 1:0.6, followed by two washes using freshly prepared 70% ethanol and resuspension in 10mM ph 8.0 Tris-HCl. 1 μg HMW-gDNA was used for nanopore library preparation and sequencing according to the standard protocol of ONT Ligation Sequencing Kit (Nanopore, https://store.nanoporetech.com/eu/ligation-sequencing-kit110.html). Sequencing was performed an ONT MinION Mk1B with three R9.4.1 flow cells.

### Re-assembly of genomes assisted by Nanopore and Hi-C

Base-calling was done by ONT Guppy v6.5.7 (--trim_adapters –trim_primers)(Wick et al., 2019). A *de novo* draft male genome assembly was generated based on Ec32 ONT data by the Canu assembler v2.2(genomeSize=220m -pacbio-raw)(Koren et al., 2017), with three iterations of error correction by Pilon v1.24 (Walker et al., 2014). An additional scaffolding step was accomplished by ARCS v1.2.5 ( z=1500 m=8-10000 s=70 c=3 l=3 a=0.3)(Yeo et al., 2018). As long read sequencing input the original ONT read data was extended by the previous assembly(Cock et al., 2010); the same strategy was also used for the new *Ectocarpus* female draft genome with Ec25 but only used ONT data.

The Hi-C raw reads underwent a preprocessing step using Trimmomatic v.0.39 with a default setting to remove the adapters and other Illumina-specific sequences ^77^. Subsequently, the clean reads were aligned draft genomes using 3D *de novo* assembly (3D-DNA) pipeline, following ^78^. The resulting Hi-C contact map, based on the initial chromosomal assembly, was visualized using Juicebox ^79^. Juicebox also facilitated the manual adjustment of contig orientations and order along the chromosomes, based on the observed contacts. During this adjustment process, some incorrectly placed sequences were trimmed from the original contigs and reassembled with the appropriate ones. The orientation of the final chromosome name was corrected with the previous reference genome ^28^. To refine the assembly, we employed TGS-GapCloser with error correction by racon v1.4.3, along with RFfiller utiliz-ing ONT reads for gap filling (Midekso & Yi, 2022; Xu et al., 2020). Subsequently, an assessment of genome quality was conducted by Benchmarking Universal Single-Copy Orthologs (BUSCO) ^32^ together with its eukaryote and stramenopiles databases in the version odb10.

To be consistent with the V2 genome, we extracted the gapless 1.55 Mb female sex determining region (SDR) of the female assembly and added it as a separate contig to the male genome (fSDR). This ‘reference’ assembly is the new *Ectocarpus sp*.7 V5 reference genome.

To identify bacterial contamination in the genome assemblies, the new assembled scaffolds were analyzed by kraken2 (version 2.1.3) ^80^, blastn (version 2.13.0, nt database 2022-07-01) ^81^ and blobtools (verson 1.1.1)^82^. Hits identified by all three tools were considered, and corresponding contamination scaffolds were removed. During the contamination analyses we removed two Hi-C scaffolds corresponding to the bacteria genera *Paraglaciecola* and *Halomonas*.

### Hi-C data analysis

The Hi-C reads were processed using the Juicer pipeline (Dudchenko et al., 2017), and binning was performed at various sizes, including 2kb, 5kb, 10kb, 20kb, 50kb, 100kb, and 500kb. The clean Hi-C data was mapped to its corresponding re-assembled reference genome (male or female *Ectocarpus* V5, Figure S1) using Bow-tie2(Langmead & Salzberg, 2013). During the alignment, the clean reads were aligned end-to-end, and spanning ligation junctions were trimmed at their 3’-end and realigned to *Ectocarpus* newly assembled genome. The result-ing aligned reads from both fragment mates were then paired and stored in a paired-end BAM file. Invalid Hi-C reads including discarding dangling-end reads, same-fragment reads, self-circled reads and self-ligation reads were removed from further analyses.

### Chromosomal contact probability

The reads information processed by the Juicer pipeline in the “merged_nodups.txt” were converted to pairs using the pairix tool (Lee et al., 2022). The draft genome was divided into 1,000 bp bins, and the contact probability *P*(s) was calculated and visualized using cooltools ^83^ following the guidelines provided in the documentation at https://cooltools.readthedocs.io/en/latest/notebooks/contacts_vs_distance.html. In short, P(s) was determined by dividing the number of observed interactions within each bin by the total number of possible pairs.

### A/B compartment identification

The A/B compartment status was determined using Eigen values (E1) obtained through eigenvector decomposi-tion of Hi-C contact maps. To calculate the E1 values at a 10kb resolution, Cooltools software was utilized with the “cooltools eigs-trans” function and GC density file ^83^. The resulting E1 values were then loaded into the plaid pattern of Hi-C contact maps. Manual validation based on intra or inter-chromosomal interactions in Hi-C was performed along each chromosome to obtain the final list of “E1” values. Since the direction of eigenvalues is arbitrary, positive values were assigned the label “A”, while negative values were assigned the label “B” based on their association with GC or gene density. The compartment border was defined as the edge bin separating the A and B compartments.

### ChIP-seq and RNA-seq

ChIP-seq and RNA-seq data from the male (Ec561) and female (Ec560) strains were obtained from ^42^. The datasets include two replicates of H3K4me3, H3K9ac, H3K27ac, H3K36me3, and H3K79me2 samples, as well as two control samples (an input control corresponding to sonicated DNA and anti-histone H3). To process the data, the nf-core ChIP-seq pipeline v2.0.0 was employed ^84^. Briefly, the raw data underwent trimming using Trim Galore v0.6.4 (Krueger, 2015), and the paired-end reads were aligned to the reference genome using BWA v0.7.17(Li & Durbin, 2009). Subsequently, MACS2 with default parameters was used to call broad and narrow peaks (Gaspar, 2018). Three replicates of RNA-seq data were trimmed by Trimmomatic v0.39 and mapped on *Ectocarpus* V5 reference genome (Figure S1) by GSNAP aligner v2021-12-17(Bolger et al., 2014; Wu et al., 2016), unique mapped read pairs were used to calculate read counts per gene by featureCount v2.0.3, DEseq2 (v1.41.6, Bioconductor) was used for detection differential expression genes with the threshold of adjusted *p* value =< 0.01 and log2fold change >= 1, TPM (Transcripts Per Million) was used for gene expression quantification (Gueno et al., 2022; Liao et al., 2014; Love et al., 2017).

### Centromere characterisation

Broad centromeric regions were determined by visually assessing the Hi-C contact maps. To assess the repeat content of these regions, RepeatModeler v2.0.2 (Flynn et al. 2020) was run on the male Ec32 V5 genome assembly to generate de novo repeat consensus models, using the flag “-LTRStruct” to perform LTR structural searches. The subsequent repeat library was provided as input to RepeatMasker v4.0.9 (https://www.repeatmasker.org/Re-peatMasker/) to identify genomic coordinates of repeats. Tandem Repeats Finder v4.09.1 (Benson 1999) was run to identify coordinates of satellite and microsatellite DNA using the recommended parameters “2 5 7 80 10 50 2000”, enabling satellite DNA with monomers up to 2 kb to be identified. Final tandem repeat coordinates were achieved by combining the simple and low complexity repeats identified by RepeatMasker with the repeats iden-tified by Tandem Repeats Finder. All other repetitive coordinates identified by RepeatMasker that did not overlap tandem repeats were assumed to be interspersed repeats (i.e. transposable elements).

Putative centromeric repeats were identified by searching for repeat families that were both almost exclusively present in the broad centromeric regions defined by the contact maps and common to all chromosomes. The two repeat models that met these criteria were then manually curated following Goubert et al. (2022). Retrotrans-posons related to *ECR-1* were identified by passing the predicted protein to Repbase Censor online tool (Kohany et al. 2006). Centromeric coordinates were defined as the first to the last copy of *ECR* elements (see Table S9). All centromeric analyses were performed on the male Ec32 V5 genome, except for the U chromosome which was analysed using the female Ec25 genome.

## Acknowledgements

This work was supported by the MPG, the ERC (grant n. 864038), the Moore Foundation (GBMF11489) and the Bettencourt-Schuller Foundation. RC is supported by a grant HORIZON-MSCA-2022-PF-01 (Project ID: 101109906). We thank the BMBF-funded de.NBI Cloud within the German Network for Bioinformatics Infrastruc-ture (de.NBI) (031A532B, 031A533A, 031A533B, 031A534A, 031A535A, 031A537A, 031A537B, 031A537C, 031A537D, 031A538A). We thank Remy Luthringer and Andrea Belkacemi for assistance with the algal cultures.

## Conflict of Interests

The authors declare no conflict of interests.

## Data accessibility

Data is available in NCBI under the project number PRJNA1105946.

## Author contributions

PL: Investigation (lead); Formal analysis (lead); Visualization (lead), Writing – original draft (equal) JV: Investigation (equal); Methodology (supporting); Formal analysis (supporting)

RC: Investigation (equal), Methodology (supporting); Visualization (equal); Formal analysis (equal); Writing – re-view and editing (supporting)

JBR: Investigation (equal), Methodology (supporting); Visualization (equal); Formal analysis (equal)

EA, CM: Investigation (supporting) MB: Methodology (supporting)

FBH, CL: Data curation (equal); Visualization (supporting); Formal analysis (equal); supervision (equal); Writing – review and editing (supporting)

SMC: Conceptualization (lead); Funding acquisition (lead); Methodology (equal); Project administration (lead); Supervision (lead); Visualization (supporting); Writing – original draft (equal); Writing – review and editing (lead).

## Notes

### Competing Interest Statement

The authors have declared no competing interest.

